# KG-Orchestra: An Open-Source Multi-Agent Framework for Evidence-Based Biomedical Knowledge Graphs Enrichment

**DOI:** 10.64898/2026.02.18.706536

**Authors:** Ahmed Hossameldin Mohamed, Karim S Shalaby, Abish Kaladharan, Heval Atas Güvenilir, Alpha Tom Kodamullil

**Author notes:** **Corresponding Author:** Alpha Tom Kodamullil, Department of Bioinformatics, Fraunhofer Institute for Algorithms and Scientific Computing (SCAI), Sankt Augustin 53757, Germany. Telephone details.

## Abstract

Biomedical Knowledge Graphs (BKGs) offer integrative representations of complex biology, yet their utility is compromised by the limitations of current construction methods: manual curation offers high fidelity but is unscalable, whereas purely automated Large Language Model (LLM) approaches often yield broad networks lacking mechanistic granularity. We present KG-Orchestra, an open-source multi-agent framework designed to build specialized, directional, cause-and-effect BKGs by enriching seed graphs. The framework focuses on increasing granularity within specific topics by leveraging Retrieval-Augmented Generation (RAG) to autonomously acquire, validate, and integrate evidence. The system orchestrates specialized agents for retrieval, schema alignment, and triplet validation with explicit, traceable provenance, transforming sparse seeds into dense, high-resolution resources. We evaluated KG-Orchestra on two specialized contexts—the mechanistic link between Nelivaptan and Alzheimer’s Disease (NADKG) and the complex probiotic interactions within the gut–brain axis (ProPreSyn-GBA)—across varying computational budgets. Our benchmarking results demonstrate that Qwen 3 variants deliver superior reasoning performance and that hybrid retrieval strategies significantly enhance evidence relevance. Furthermore, the multi-agent architecture ensures high triplet integrity and biological validity through iterative cross-checking and self-correction. The framework remains computationally flexible, deploying from single laptop GPUs to high-performance clusters. By bridging knowledge gaps and adding context-aware entities, KG-Orchestra increases reliability while validating seed assertions against up-to-date sources. This versatility supports critical downstream applications, including completing missing mechanistic pathways, integrating novel entities for drug repurposing, constructing targeted subgraphs from entity lists, and retroactively validating graph evidence for transparent auditing.

## 3. Introduction

The digitization of Electronic Health Records (EHRs), coupled with the rapid democratization of high-throughput omics technologies and the proliferation of wearable sensing devices, has fundamentally altered the medical information landscape [1]. While this exponential growth of data has driven numerous innovations in healthcare [2], it has also created a significant challenge: information fragmentation [3]. Parallel to the expansion of structured clinical data, there is a massive, ever-growing body of scientific publications containing critical but "hidden" unstructured insights [4]. Navigating the semantic complexities of this literature and integrating it with high-dimensional clinical data is notoriously difficult [5]. Traditional data management strategies, such as relational databases, are adept at storing isolated records but face significant limitations in capturing the intricate, multidimensional relationships that define biological systems [6]. Consequently, there is a critical need for frameworks that can effectively distill these unstructured insights into a human-readable and machine-computable form, enabling the construction of highly granular representations of complex biological processes.

Biomedical Knowledge Graphs (BKGs) organize complex domain knowledge into structured networks where nodes represent biological entities—such as genes, drugs, diseases, and proteins—and edges define the semantic relationships between them (e.g., Protein X activates Gene Y or Drug A treats Disease B). By explicitly mapping these interactions, BKGs integrate multimodal and multiscale data into a unified framework. This structured representation provides a vital strategy for overcoming traditional data management strategies limitations, offering a more detailed and holistic understanding of biological systems [7]. With this, BKGs support sophisticated tasks of modeling diseases, thereby supporting identification of therapeutic targets, drug repurposing, and personalized treatments — capabilities that traditional data systems often lack [6]. Advances in artificial intelligence, especially in large language models (LLMs), have played a key role in enhancing the design and usefulness of these graphs [8]. Approaches like graph-based machine learning and language processing tools have extended the range of applications, allowing researchers to uncover hidden patterns in disease biology and speed up drug development pipelines, and demonstrating how BKG can serve as powerful tools for advancing biomedical science by offering an integrated and flexible way to study intricate biological systems [6].

The construction of BKGs involves a multi-step pipeline starting with data acquisition from structured databases, semi-structured repositories, and unstructured text [6]. After preprocessing for noise reduction and normalization, relationships are extracted and represented as triples (Head–Relationship–Tail). Integration then aligns these entities and relations across datasets to form a unified graph [9–11]. These workflows can be performed manually by expert curators to ensure high precision, or automatically using computational methods to address the challenge of data scalability. Early automated approaches primarily relied on supervised Natural Language Processing (NLP) pipelines, utilizing specific architectures such as Convolutional Neural Networks (CNNs) or Recurrent Neural Networks (RNNs) for Named Entity Recognition (NER) and Relation Extraction (RE) [12–14]. Several large-scale, widely used BKGs have been constructed by integrating multiple databases into a single heterogeneous network, including PrimeKG (integrating 20 resources across molecular, clinical, and pharmacological scales) [15], Hetionet (assembled from 29 databases) [16], and SPOKE (connecting information from 41 databases) [17]. In parallel, other resources are generated fully or partially through automated extraction from the biomedical literature to improve scalability and coverage, such as SemMedDB [18], GNBR [19], and assembly frameworks like INDRA [20] that consolidate outputs from multiple text-mining systems. While these integrated and literature-derived resources provide valuable breadth and a practical substrate for downstream machine learning, they are often insufficient as a sole evidence base for mechanistic or decision-support applications due to heterogeneous evidence quality, normalization and mapping errors, a substantial fraction of associative rather than causal relationships, and version lag that can limit alignment with the rapidly evolving biomedical literature. In automatically extracted BKGs, additional challenges arise from limited semantic interpretation, loss of experimental context, and insufficient evidence traceability. Consequently, the reliability and flexibility of the underlying extraction workflows remain critical considerations. While effective within closed domains, these traditional methods are often limited by their reliance on extensive labeled datasets and rigid schemas, which hinders their adaptability to new entity types [21]. Recently, the field has shifted toward Large Language Models (LLMs), which offer significant advantages over traditional NLP approaches by replacing discriminative classification with generative extraction [8]. Unlike their predecessors, LLMs leverage massive pre-training to perform few-shot or zero-shot learning, allowing them to extract complex relationships from low-resource domains without the need for task-specific fine-tuning [22]. Furthermore, LLMs demonstrate superior semantic reasoning capabilities, enabling them to capture implicit or long-range dependencies in text and handle open-world information extraction more robustly than rule-based or pipeline-based systems [23,24].

Recent applications have leveraged LLMs to automate the construction of KGs by extracting triplets from scientific papers; for instance, research efforts by Wu et al. [25] and Lee et al. [26] have explored using LLMs to extract entities and relationships at scale. Another notable work by Arsenyan and Bughdaryan demonstrated an end-to-end framework for extracting relations from electronic medical records (EMRs), successfully constructing a KG for age-related macular degeneration that interlinked treatments, risk factors, and symptoms [27]. Meanwhile, the ACKG-LLM framework introduced a modular design that divides the construction process into subtasks, i.e., information extraction, relational semantic enhancement, and schema normalization, to reduce manual effort [28]. To address the limitations of monolithic models in handling such complex workflows, the field is increasingly adopting Multi-Agent Systems (MAS). Unlike single-agent architectures, MAS decompose intricate tasks into manageable sub-components assigned to specialized agents, thereby enhancing solvability through divide-and-conquer strategies [29]. This collaborative paradigm significantly outperforms isolated agents by enabling iterative cross-verification and debate, which reduces hallucination rates and improves reasoning accuracy in knowledge-intensive tasks [30,31]. Furthermore, assigning distinct roles—such as critic, generator, and verifier—allows these systems to maintain context and consistency over long-horizon tasks better than a single LLM attempting to manage the entire extraction pipeline simultaneously [32]. Building on these construction and multi-agent capabilities, automated enrichment of BKGs has recently gained traction with frameworks like KARMA, which utilizes a multi-agent, LLM-driven architecture to identify novel entities and resolve conflicts within evolving literature [33]. However, despite the success of such multi-agent systems, they remain heavily dependent on the coverage of their source corpora; incomplete or biased collections inevitably propagate gaps into the resulting KG, limiting the ability to capture the dynamic and granular contradictions inherent in biomedical discovery.

Despite advances in automation, biomedical research still relies on unscalable manual curation to ensure data quality, as automated methods often lack the necessary precision. To overcome this bottleneck and the biases inherent in static literature corpora, there is a critical need for frameworks that combine automated scalability with high-fidelity extraction. By moving beyond simple verification to capture highly granular, causal relationships, such enriched graphs allow researchers to understand the mechanistic "why" behind biological phenomena rather than merely observing correlations, ultimately empowering deeper and more precise scientific discovery.

Motivated by the critical need to move beyond static, general-purpose graphs toward highly granular, context-specific representations, we propose KG-Orchestra, an open-source multi-agent framework designed to autonomously enrich seed-knowledge graphs, which are high-quality BKGs constructed manually by extracting knowledge from limited number of scientific articles, into specialized, high-resolution discovery engines, by acquiring, assessing, and integrating domain-specific biomedical knowledge based on evidence lines from scientific publications. KG-Orchestra combines retrieval-augmented generation with specialized generative agents to proactively assemble relevant literature and data, cross-validate current and newly extracted triples against source and external evidence, and harmonize entities and relations with the target schema, enabling scalable and seamless enrichment without reliance on limited precompiled corpora. By adding missing knowledge to seed graphs and transforming them into dense, highly granular, and biologically valid networks, KG-Orchestra enables researchers to interrogate the mechanistic knowledge of specific domains—linking entities through explicit, evidence-backed pathways. This approach advances biomedical research not merely by increasing data volume, but by delivering the depth and causal clarity required to uncover latent connections, validate novel hypotheses, and drive truly knowledge-driven innovation.

## 4. Methods

The following sections describe the KG-Orchestra framework, its multi-agent architecture, and the procedures used for knowledge graph validation and enrichment, and the framework validation. All components of the system — including orchestration logic, agent specifications, prompt templates, and enrichment workflows — are available in the accompanying open-source repository for reproducibility (GitHub: https://github.com/Fraunhofer-SCAI-Applied-Semantics/KG-Orchestra).

### 4.1 Framework Overview

KG-Orchestra is a multi-agent LLM framework for validating and expanding biomedical knowledge graphs by extracting entities and relations from curated corpora and web sources. As illustrated in Figure 1, starting from a seed KG of typed head–relation–tail triples with evidence, it iteratively validates each triple and seeks indirect, directional paths connecting head to tail. The workflow formulates a templated query, retrieves candidate paragraphs via a vector-store pipeline with an LLM Paragraph Evaluator, and adjudicates seed triples as valid or need-review with full provenance. An LLM Path Builder then constructs directed single- or multi-hop mechanistic/functional/causal paths, which undergo triplet-level processing: schema alignment (with controlled schema extension), entity linking/creation, evidence-augmented validation, and repair attempts when invalid. Validated triplets are integrated into the KG or enrich existing ones with PubMed Central/DOI-backed excerpts, preserving provenance. If evidence is insufficient, the system falls back to PubMed retrieval and retries; otherwise, it proceeds to the next seed triple—providing a comprehensive, auditable framework for KG validation and extension.

**Figure 1.**
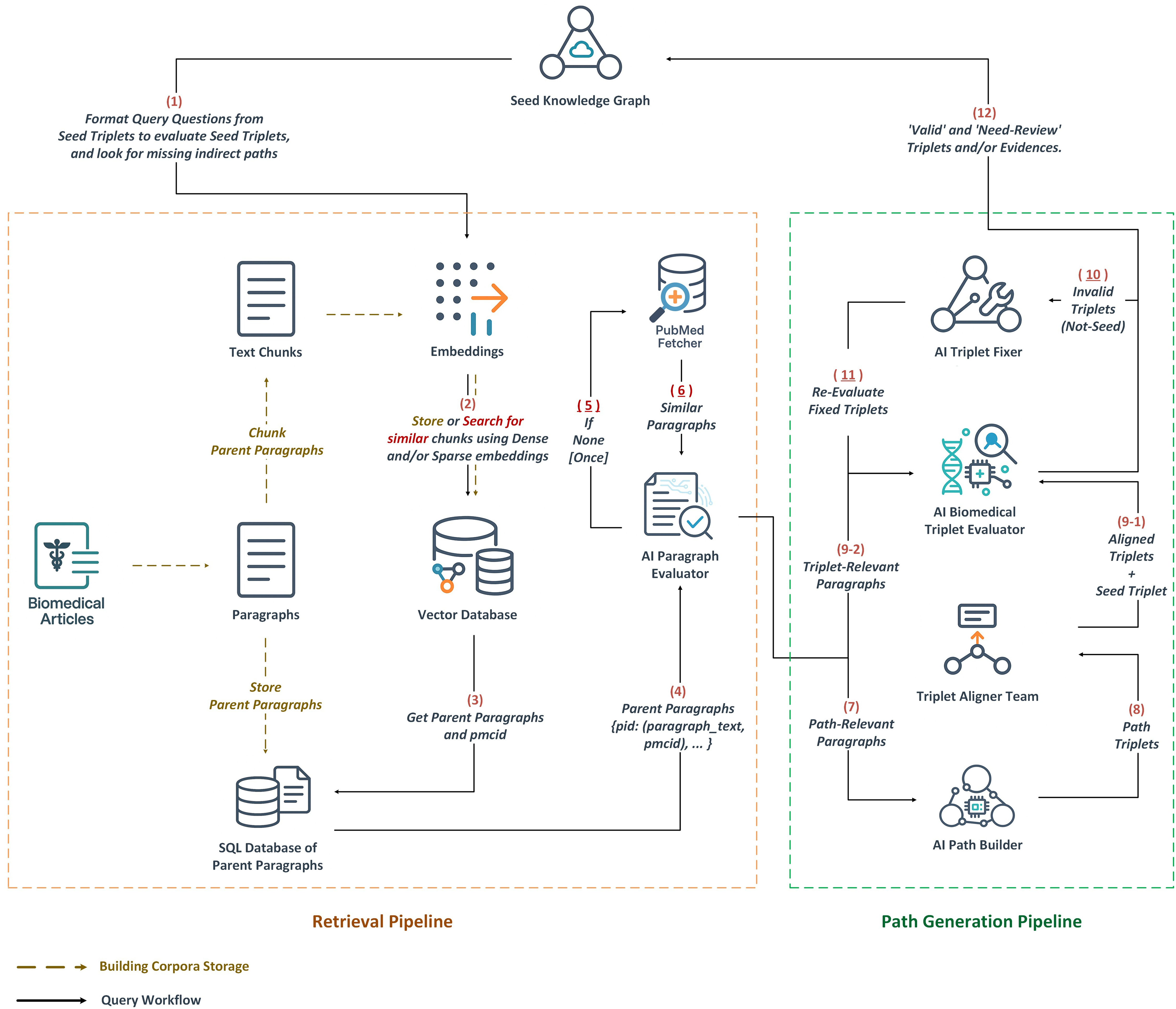
KG-Orchestra query workflow: Starting from a seed knowledge graph (Seed KG) comprising typed, directed triples T = (head, relation, tail, e), where e denotes evidence, the system iterates over selected triples to discover missing indirect paths from head to tail. For each seed triple, KG-Orchestra (1) formulates a templated question that preserves entity types and source-to-target directionality; (2-7) retrieves candidate paragraphs (parent paragraphs) from a vector store of embedded biomedical text and uses an LLM Paragraph Evaluator to retain only relevant chunks; (8) constructs a directed multi-triplet path p = [T1, T2, …, Tk] via Path Builder Agent; (9-11) conducts triplet-level processing and validation, including schema alignment (mapping or minimally extending entity and relation types) by Schema Aligner Agent, entity resolution (linking to existing nodes or creating new ones), evidence augmentation and verification (additional retrieval and LLM-based checking by Triplet Evaluator agent), and correction by Triplet Fixer agent, with need-review flagging if unresolved; (12) integrates validated triples into the graph or enriches existing ones with provenance (e.g., PubMed IDs, DOIs, supporting excerpts); and (5-6) if evidence is insufficient, falls back to PubMed search for relevant paragraphs and retries path construction, otherwise proceeding to the next seed triplet. (9-1, 9-2) original seed triplet is also evaluated by the Triplet Evaluator agent, adding additional contradicting/supporting evidence, and labeled as valid or need-review, similar to the newly introduced triplets.

### 4.2 Definition of Evidence in the KG-Orchestra Framework

Within the KG-Orchestra workflow, evidence refers to a paragraph-level textual excerpt retrieved from biomedical literature that explicitly articulates the semantic relation encoded by a candidate head–relation–tail triple. Evidence is thus operationally defined as a bounded unit of text that provides direct support for, or contradiction of, a specific triplet under evaluation or construction. These excerpts are sourced through the system’s templated query and vector-store retrieval pipeline, then assessed by the LLM Paragraph Evaluator for relevance to the target relation. Importantly, each evidence unit maintains full provenance, including publication identifiers (e.g., DOI, PubMed/PMC ID). This traceability allows for downstream appraisal of study design if needed. Evidence may accompany both seed triples undergoing validation and newly generated triples created by the LLM Path Builder, ensuring traceability and supporting auditable KG expansion. Therefore, and in alignment with the Evidence and Conclusion Ontology (ECO) [34], Evidence within KG-Orchestra workflow is formally categorized as “author statement supported by traceable reference used in automatic assertion” (ECO:0007321).

This definition is distinct from, yet compatible with, the concept of evidence in Evidence-Based Medicine (EBM). EBM classifies evidence according to study design and methodological rigor [35], whereas KG-Orchestra focuses on textual claims extracted from those studies, independent of their position in the EBM hierarchy. The framework does not assess study quality; rather, it determines whether the retrieved paragraph expresses a mechanistic, functional, or causal assertion that aligns with—or contradicts—the KG triple under review.

### 4.3 Evidence-Retrieval Pipeline

To support evidence retrieval across diseases, KG-Orchestra operates over a large, open-access biomedical text collection. As a proof of concept, we assembled a domain-focused corpus comprising the full text of 62,173 open-access biomedical articles indexed under the Medical Subject Headings (MeSH) term: “Neurodegenerative Diseases” (as of August 2025) From PubMed Central database. Corpus preparation comprised two subprocesses: (i) semantic-preserving, computationally efficient chunking and (ii) embedding-based indexing for retrieval.

#### 4.3.1 Query formulation

A templated question to preserve directionality and typing is instantiated: “What is the biomedical pathway that connects [*head name and type*] as the source to [*tail name and type*] as the target, with direction from source to target?”

#### 4.3.2 Text chunking strategy

From the outset, corpus preparation revolved around a single practical tension: the granularity at which text should be indexed [36]. Sentences are too brittle for mechanistic reading—biomedical pathways often unfold across multiple sentences—yet paragraphs can exceed encoder limits and inflate computing. Therefore, we prioritized the following chunking scheme to reserve semantics while remaining tractable at corpus scale and robust to encoder maximum sequence length limits. First, we split each paragraph into smaller, manageable units (sentence-level or 512-token-length–bounded segments) and embed these units individually into a dense/sparse vector space. Afterward, the original parent paragraphs persisted in a relational database and a mapping from each embedded unit to its parent paragraph was maintained. Retrieval returns fine-grained chunks for matching and promotes the corresponding parent paragraphs for downstream extraction and path construction. To evaluate the retrieval quality of sentence-level versus 512-token-length–bounded text chunking, we used three BEIR datasets [37], and one in-house dataset using normalized Discounted Cumulative Gain at 10 (*nDCG*@10) computed with the BEIR library (Table 1). *nDCG*@*K* measures ranking quality up to position *K* by computing discounted cumulative gain of the *topK* results and normalizing it by the ideal *nDCG*@*K*, capturing both relevance and position [38].

**Table 1.**
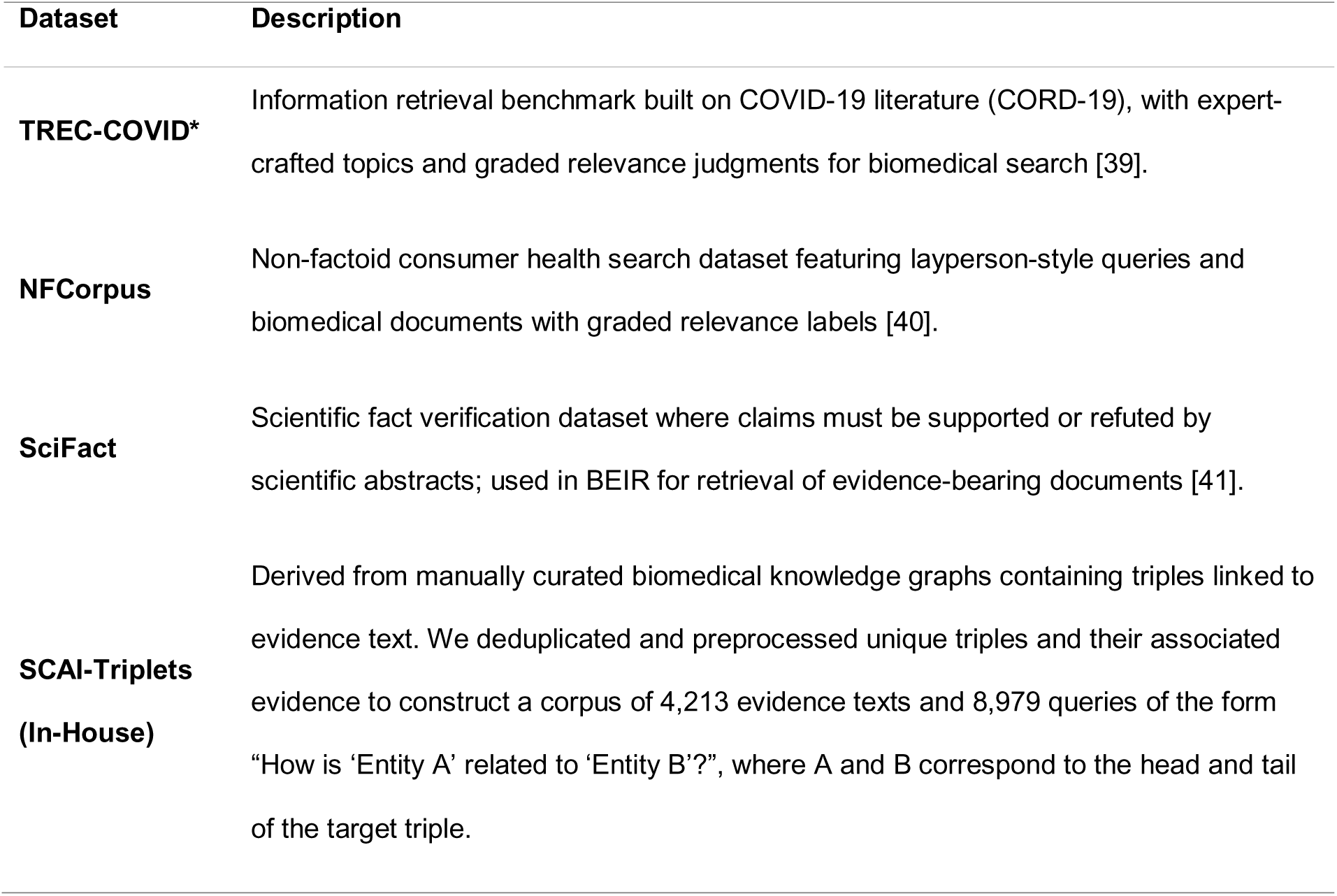
The benchmark datasets used to evaluate the embedding and retrieval along with the dense embedding models studied.

#### 4.3.3 Embedding model selection

To select the final retrieval configuration, we compared twelve dense embedding models (Table 2), across NFCorpus, SciFact, and SCAI-Triplets datasets using *nDCG*@10 . TREC-COVID wasn’t used to benchmark embedding models due to its comparatively rich corpus (171K text elements), which encounters high computational costs.

**Table 2.**
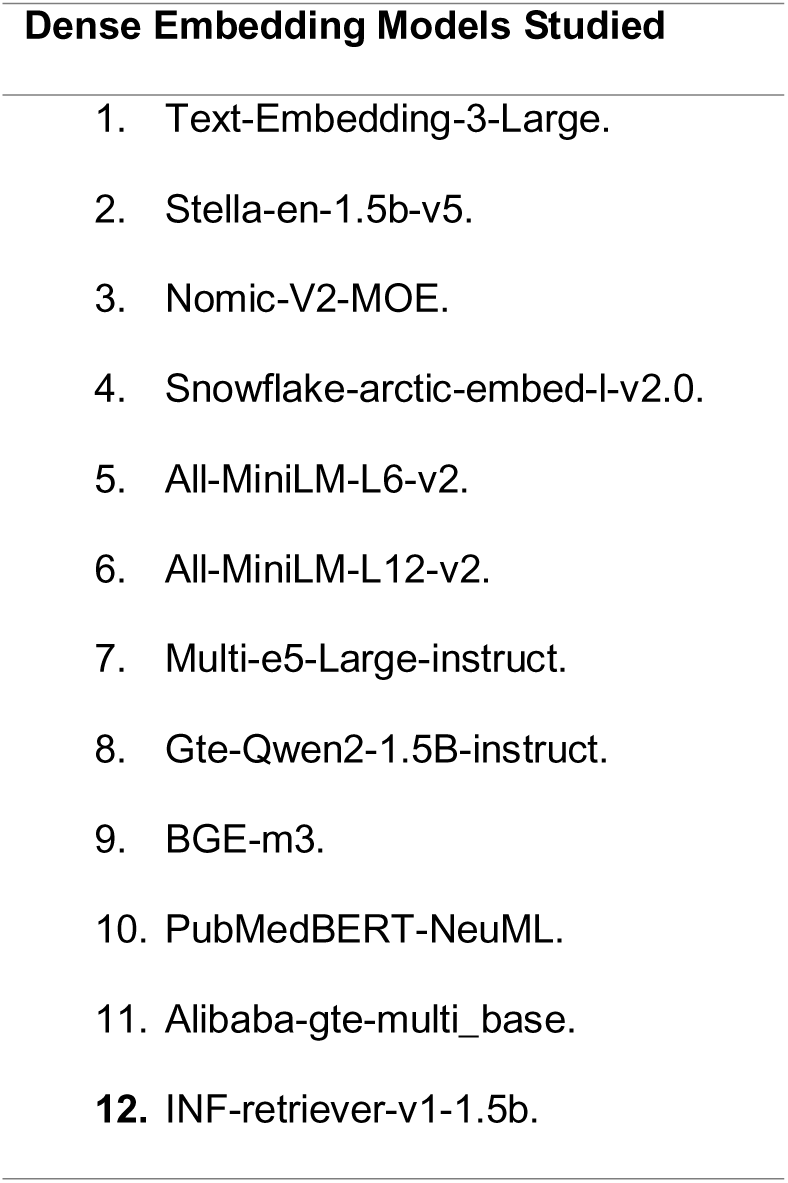
The dense embedding models studied.

#### 4.3.4 Embedding and retrieval methods

The density of the resulting vector embedding of chunks differs according to the embedding strategy and would affect the precision and robustness of retrieval. Therefore, we evaluated the dense, sparse, and hybrid retrieval to balance precision, and robustness to terminology variation.

##### 4.3.4.1 Dense embeddings

We benchmarked compact to mid-size open-source dense embedding models (both general-purpose and domain-finetuned) and, for reference, OpenAI’s text-embedding-3-large, given the strong semantic matching and context sensitivity of dense models [42].

##### 4.3.4.2 Sparse embeddings

We used SPLADE-v3, which applies a masked-LM head with sparsity-inducing regularization (e.g., L1/FLOPs) and pooling to produce interpretable, sparse term-weight vectors while expanding to semantically related terms to mitigate vocabulary mismatch [43].

##### 4.3.4.3 Hybrid search

We independently scored candidate chunks with a dense model and a SPLADE-v3 model, then fused scores using a distribution-based score fusion function in Qdrant Vector Database to obtain the final ranking [44]. Qdrant Vector Database is well-suited for open-source hybrid search because it can store multiple representations per document (dense embeddings and sparse term-weight vectors) and fuse their scores, improving ranking by combining semantic and lexical signals. Its fast ANN indexing (HNSW), filterable payloads, and self-hostable, reproducible setup make experiments easy to run, compare, and share without vendor locklZlin. The combination of dense embeddings and SPLADE-v3 representation typically improves robustness over single-method retrieval, especially under paraphrase and domain-specific terminology.

### 4.4 Biomedical Path Construction Pipeline

Following retrieval, embedded chunks are mapped back to their parent paragraphs. These paragraphs then pass through a multi-agent pipeline to extract relevant biomedical knowledge and assemble a directed path connecting the query head to the query tail.

#### 4.4.1 Path Construction

Pathway construction begins using a Paragraph Evaluator to label retrieved parent paragraphs as strongly or partially relevant so complementary evidence can be combined. The Path Builder composes a single directed chain of evidence-backed triplets; if evidence is insufficient, a PubMed Web Fetcher performs hybrid dense+sparse retrieval with score fusion to supply additional paragraphs for another build attempt. Each triplet then undergoes triplet-level processing: a Schema Aligner maps entities and relations to the seed KG schema (extending types only when necessary), an Entity Matcher resolves entities via exact match or using the Unified Medical Language System (UMLS)-based normalization (creating nodes only when needed) [45], and evidence-based validation evaluate final triplets before integration into the seed KG.

#### 4.4.2 Paragraph Evaluator

Given the query and the retrieved paragraphs, the Paragraph Evaluator assesses each paragraph’s contribution to answering the query, with the expectation that strongly and partially relevant paragraphs will be combined to recover the full path. Each paragraph is labeled as “strongly relevant” if the paragraph alone establishes a mechanistic, functional, or associational link between the head and tail, “Partially relevant” if the paragraph contributes complementary information but does not fully answer the query, or “Irrelevant” if the paragraph is unlikely to contribute useful information to path construction, even in combination.

#### 4.4.3 Path Builder

Using the set of strongly and partially relevant paragraphs, the Path Builder composes a single directed path from head to tail. The output is a linear sequence of triplets [head, relation, tail], where each triplet is a direct, evidence-backed relation (directional, mechanistic/causal/functional/conceptual) between biomedical entities.

#### 4.4.4 PubMed Web Fetcher (fallback)

If no suitable path can be constructed due to insufficient evidence or a lack of relevant paragraphs, the PubMed Web Fetcher queries PubMed for articles that mention both the head and tail in their abstracts or full texts. Available full texts (or abstracts when full text is unavailable) are segmented into paragraphs, embedded, and ranked via hybrid retrieval (distribution-based score fusion of dense and sparse models). The top-k paragraphs are then passed to the Paragraph Evaluator; strongly and partially relevant paragraphs are fed back into the Path Builder.

#### 4.4.5 Triplet-Level Processing

Each triplet produced by the Path Builder is subjected to schema alignment, entity resolution, and evidence-based validation before integration into *Seed KG*.

##### 4.4.5.1 Schema Aligner

The Schema Aligner maps triplet entities and relations to *Seed KG* schema, aligning entity types to high-level biomedical classes and relations to polarity-aware mechanistic/causal types. When no suitable entity or relation type exists, the schema is extended to introduce the required type, minimizing unnecessary proliferation while preserving granularity.

##### 4.4.5.2 Entity Matcher

Aligned triplets undergo entity resolution to map triplet entities to existing nodes and avoid redundancy. Entity resolution proceeds in three stages: first, we query the seed KG for an exact match on entity name and type; if found, the entity is mapped to that node. If no exact match exists, we normalize to UMLS by encoding the entity’s name and type and querying a vector index of UMLS concepts that combines dense and sparse (lexical) embeddings derived from concept names and types; we select the nearest concept and obtain its Concept Unique Identifier CUI, then map the entity to an existing KG node with the same CUI if present. If neither an exact match nor a CUI match is available, a new node is created.

##### 4.4.5.3 Triplet Validation Team

After entity resolution, triplets are validated, for biological plausibility, semantic coherence, and directional correctness, through several steps including:

##### 4.4.5.4 Evidence augmentation

Up to two additional paragraphs, explaining the head–tail relationship, are retrieved via the Evidence-Retrieval Pipeline.

##### 4.4.5.5 Triplet evaluation

A Triplet Evaluator reviews the triplet against the original parent paragraph and augmented evidence(s), labeling the triplet as valid or invalid based on:

**a. Biological validity:** The relation is biologically plausible and supported by evidence; entity names are unambiguous and correctly typed.
**b. Semantic coherence:** The triplet structure and relation are used appropriately in context.
**c. Directionality:** The head-to-tail direction is biologically and logically sound.

##### 4.4.5.6 Triplet repair

Invalid triplets are passed to a Triplet Fixer that attempts to correct the relation and triggers re-evaluation. If repair fails, the triplet is labeled need-review. Both valid and need-review triplets are accompanied by evaluator-generated rationales explaining the decision.

#### 4.4.6 Graph Integration and Provenance

Lastly, validated and need-review triplets, together with their originating and augmented evidence (parent paragraph, PubMed IDs/DOIs, and supporting excerpts), are added to *Seed KG* to form the enriched KG. Need-review triplets are flagged for subsequent manual curation.

### 4.5 Backbone LLM Selection and Agent Implementation

To identify the backbone LLM for our agent-based enrichment framework, we benchmarked four open-source, moderate-size models (≤32B parameters; Table 3) by enriching a manually curated KG (NADKG-N100 from Methods section 8.1). All experiments were conducted on a single NVIDIA A100 (40 GB) [46] using uniform 4-bit weight quantization to satisfy memory constraints. Agents were orchestrated with LangChain [47] and served via Ollama [48]. The model demonstrating the best enrichment quality under these constraints was selected for full-graph deployment. After selecting the KGlZlOrchestra’s backbone LLM, we examined scaling effects on enrichment quality and coverage by comparing variants of different sizes (number of parameters) under identical prompting and evaluation settings.

**Table 3.**
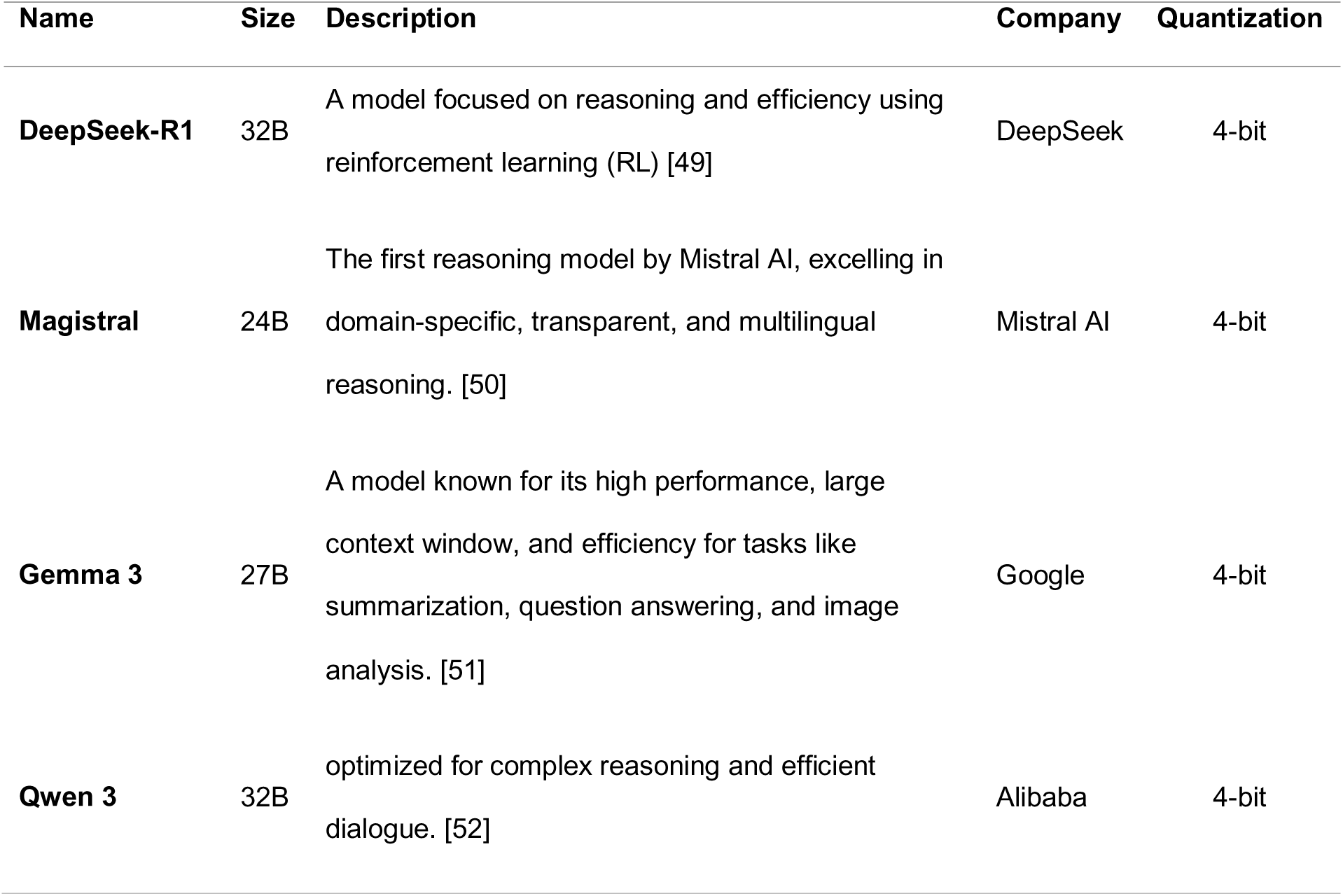
Candidate LLMs, for backbone LLM selection and agent implementation, and their information.

### 4.6 Evaluation Metrics and Protocol

We evaluated the final triplets integrated into the seed knowledge graph using a two-tier approach; automated bulk evaluation with an OpenAI GPT-5–based evaluator [53], which was used to evaluate the output triplets in all studies that lead to the structure of Kg-orchestra, and human expert evaluation on a random sample of the final output by experienced curators. Each path was assessed at both the path-level and the triplet-level labeled True/False for the criteria in Table 4, following deterministic rules.

**Table 4.**
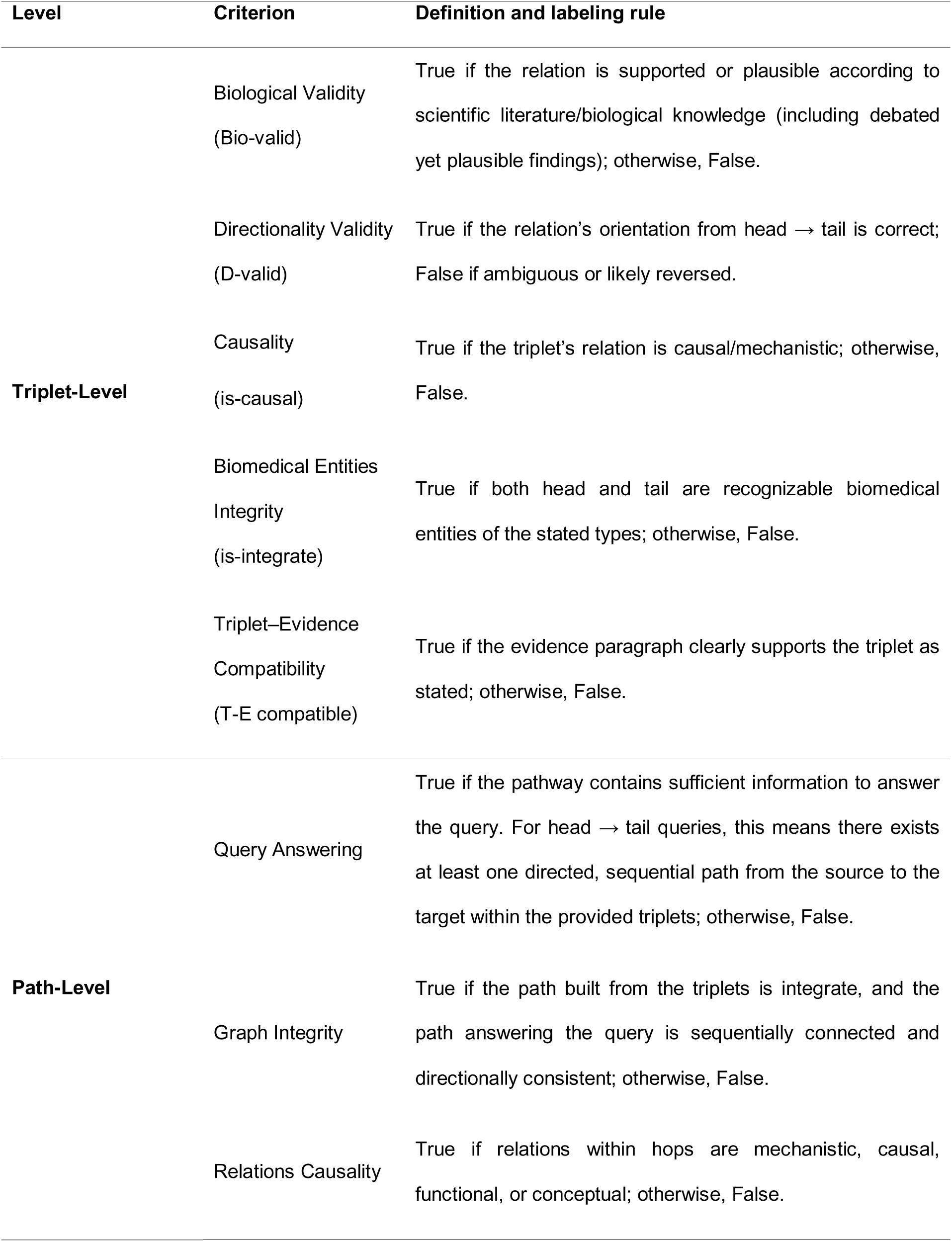
Triplet-level and Path-level evaluation criteria.

### 4.7 Ablation Study

We assessed each agent’s contribution to triplet-level quality by sequentially evaluating the generated triplets at each stage of pathway construction and validation, using the previously defined metrics. To quantify the effect of the Schema Aligner, we also executed the enrichment pipeline on a seed KG (See Methods Section 8) with this agent disabled and measured changes in triplet-level quality and in the resulting counts of entity and relation types.

### 4.8 Use case scenarios

We evaluated KG-Orchestra by enriching two manually curated biomedical seed knowledge graphs to simulate real-world deployment. The Nelivaptan–Alzheimer’s Disease KG (NADKG) investigates the understudied connection between Nelivaptan (a V1b receptor antagonist) and Alzheimer’s disease symptom pathways, while the ProPreSyn-GBA KG encodes how probiotics, prebiotics, and synbiotics modulate the gut–brain axis.

#### 4.8.1 NAD Knowledge Graph

To assess performance in real-world settings, we evaluated the proposed enrichment framework on a biomedical seed knowledge graph, manually curated from twenty scientific articles. To keep the granularity and domain focus of the seed KG, we set up a Nelivaptan–Alzheimer’s Disease KG (NADKG), that is centered on the potential connection between Nelivaptan (antagonist of V1b receptors, used in major depressive disorder) [54] and Alzheimer’s disease, with the goal of identifying pathways relevant to Alzheimer’s symptom management. The connection between this drug and AD is not well studied, so we selected this scientific gap to test if KG-orchestra can collect more granular and dense knowledge about it. Beyond full-graph, end-to-end enrichment evaluation, we generated two pruned versions of the NADKG to assess model performance under varying graph sizes. Specifically, the original NADKG (1685 nodes, 3273 triplets, 28 relation types, 21 node types) was systematically reduced, by selecting 6 and 9 out of the 20 papers as the source for the BKG construction, to obtain subgraphs containing 100 (NADKG-N100; 83 nodes, 100 triplets, 5 relation types, 8 node types) and 362 triplets (NADKG-N362; 273 nodes, 362 triplets, 6 relation types, 10 node types), respectively. These pruned graphs were used to benchmark LLMs for agent-backbone selection and to quantify the contributions of individual agent components through ablation. In addition, the two subgraph sizes enabled controlled evaluation of how initial graph scale influences downstream knowledge quality and coverage.

#### 4.8.2 ProPreSyn-GBA Knowledge Graph

As an additional real-world evaluation, we applied the proposed framework to enrich a seed knowledge graph (KG) derived from the peer-reviewed review article “A Comprehensive Overview of the Effects of Probiotics, Prebiotics, and Synbiotics on the Gut–Brain Axis” (ProPreSyn-GBA) [55]. From a biological perspective, elucidating these effects is critical because probiotics, prebiotics, and synbiotics modulate the gut–brain axis—a complex bidirectional network integrating neural, endocrine, immune, and metabolic pathways. By gathering more granular knowledge about this network like microbiota composition and activity, these interventions offer potential benefits for neurocognitive performance, emotional regulation, and immune homeostasis. Furthermore, newly added entities and relations contribute to better understanding of their role in neurodegenerative and neuropsychiatric disorders, as well as in the management of chronic inflammatory conditions. Ultimately, mapping these interactions is essential to address current gaps in mechanistic understanding and long-term clinical relevance. ProPreSyn-GBA has been constructed manually by professional curators and had 531 nodes, 1093 triplets, 6 relation types, and 10 node types.

### 4.9 Reproducibility Study

We assessed the reproducibility of KG-Orchestra by executing the enrichment pipeline on NADKG-N362 three times under identical conditions (Qwen3-32B model; NVIDIA H100, 95-GB GPU). To quantify run-to-run consistency, we evaluated the semantic similarity of the triples generated in each run using an embedding-based approach rather than exact string matching. Each triple was encoded using KaLM-Embedding-Gemma3-12B-2511 embedding model, which achieves state-of-the-art performance in Massive Multilingual Text Embedding Benchmark (due to 11-2025; see MTEB Leaderboard at https://huggingface.co/spaces/mteb/leaderboard, accessed 28 Nov 2025) [56], and pairwise cosine-similarity matrices were computed between the triples of each pair of runs. For each run pair, we quantified similarity using Best Match Average similarity [57]:

Let 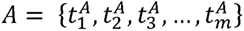 and 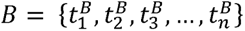 denote the sets of triplets produced by Run *A* and Run *B*, respectively, and 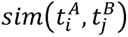 is the pairwise cosine similarity of triplet embeddings. We compute a symmetric, set-level semantic similarity between *A* and *B* by the following procedure:

1- **Best-match scores from *A*** to ***B***: For each triplet 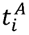 ∊ *A* compute its best-match similarity in *B*:

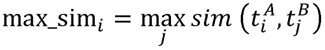

The directional mean similarity from *A* to *B* is the average of these maxima:

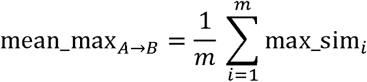

2- **Best-match scores from *B* to *A*:** Similarly, compute the directional mean similarity from *B* to *A*:

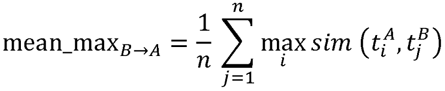

3- **Symmetric set-level similarity:** The final semantic similarity between the two runs is the arithmetic mean of the two directional scores:

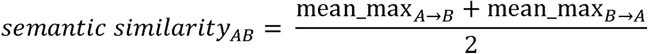

We additionally summarized key graph features—numbers of triples, nodes, entity types, and relation types—across runs. Together, these analyses provide both relation-dependent and relation-agnostic measures of the semantic reproducibility and stability of seed-KG enrichment under fixed computational and model settings.

### 4.10 Effect of Seed KG Size on Knowledge Coverage and Quality

To evaluate how the size of the initial Seed KG influences downstream knowledge retrieval and triplet quality, we compared three versions of the NADKG with increasing scale: the pruned graph containing 100 triplets (NADKG-N100), the medium-sized version containing 362 triplets (NADKG-N362), and the full NADKG (having 2516 triplets). All three graphs were subjected to the same enrichment workflow, after which we assessed their structural properties and triplet-level semantic quality. This multi-scale analysis enables quantification of how initial graph size affects overall knowledge coverage and the fidelity of generated triplets.

## 5. Results

In this section, we present a systematic evaluation of the KG-Orchestra framework, organized into three distinct phases. We begin by analyzing the fundamental parameters governing information retrieval, comparing various embedding models and hybrid retrieval strategies to optimize evidence extraction from biomedical literature. Subsequently, we conduct a comprehensive ablation study to isolate and quantify the specific contributions of each agent within the multi-agent architecture, ensuring that every component adds measurable value to the workflow. Finally, we demonstrate the system’s holistic performance and biological validity through two real-world case studies: the construction of the Nelivaptan–Alzheimer’s Disease (NADKG) graph and the enrichment of the ProPreSyn-GBA network.

### 5.1 Token-Length–Bounded Chunking Enhances Retrieval Relevance

Across four benchmark corpora (TREClZlCOVID, NFCorpus, SciFact, and SCAI-Triplets), TokenlZllength–bounded Hybrid Chunking consistently achieved higher *nDCG*@10 than SentencelZlbased Hybrid Chunking (Figure 2). On TREClZlCOVID, *nDCG*@10 increased from 0.598 to 0.774 (+0.176); on NFCorpus, from 0.304 to 0.370 (+0.066); on SciFact, from 0.561 to 0.748 (+0.187); and on SCAI-Triplets, from 0.473 to 0.498 (+0.025). While the improvement on SCAI-Triplets was marginal, since the dataset corpora chunks are mostly sentence-based, Results on TREClZlCOVID, NFCorpus, and SciFact indicate that token-lengthlZlbounded chunks are more effective for relevantlZlchunk retrieval than sentencelZllevel chunks, making the tokenlZllength–bounded scheme the preferred choice among the two strategies evaluated.

**Figure 2.**
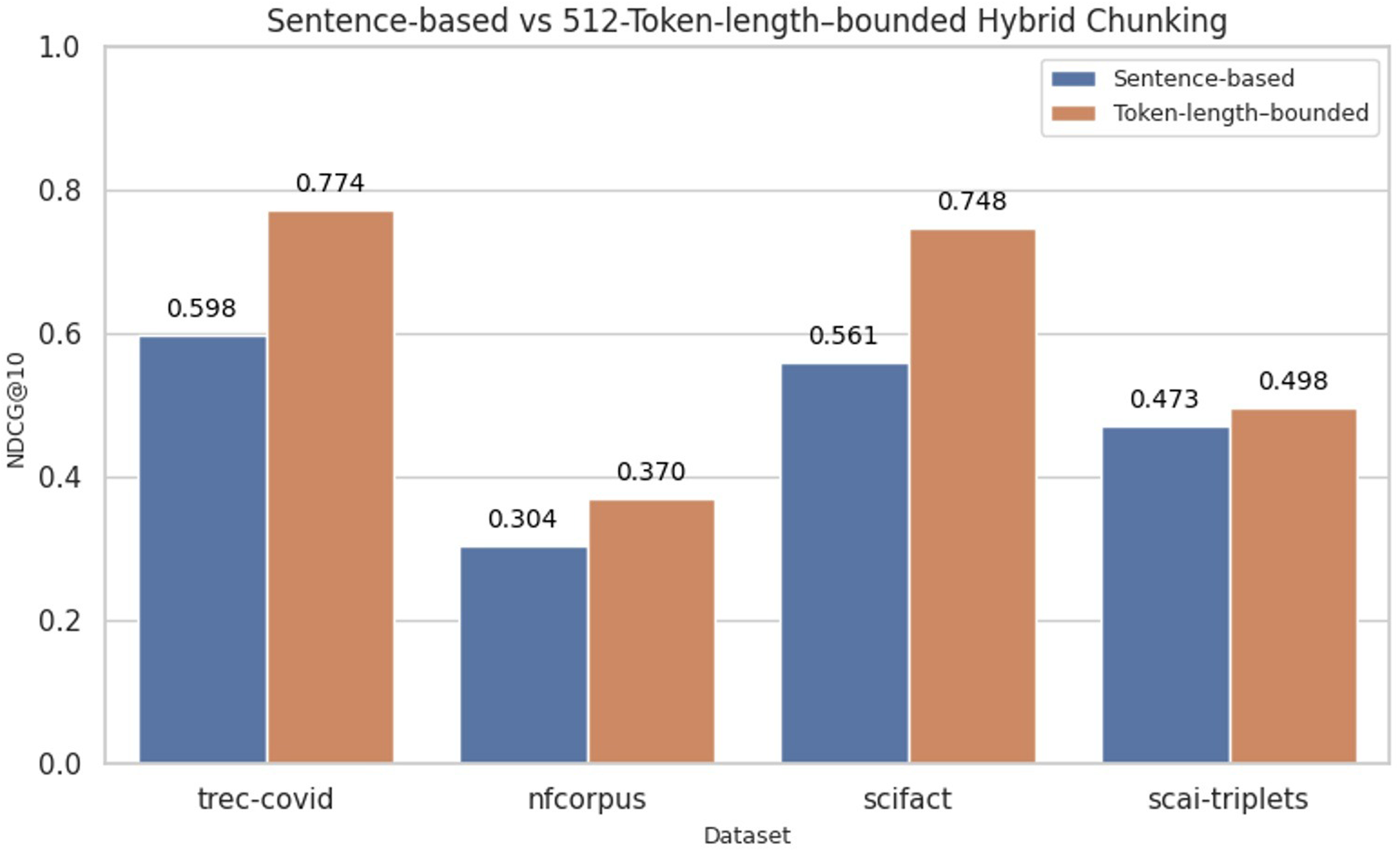
Comparison of SentencelZlbased versus TokenlZllength–bounded Hybrid Chunking across TREClZlCOVID, NFCorpus, SciFact, and SCAI-Triplets corpora.

### 5.2 Nomic-V2-MOE Delivers Superior Computational Efficiency

Across NFCorpus, SciFact, and SCAIlZlTriplets (Figures 3), *nDCG*@10 varied by model and dataset. OpenAI’s textlZlembeddinglZl3lZllarge achieved the highest scores on NFCorpus (0.419) and SCAIlZlTriplets (0.511), while StellalZlenlZl1.5BlZlv5 led on SciFact (0.777). Among openlZlsource models, StellalZlenlZl1.5BlZlv5 (1536lZld) and NomiclZlV2lZlMOE (768lZld) formed the top tier, with Nomic trailing Stella by 0.031 on NFCorpus (0.397 vs 0.366), 0.038 on SciFact (0.777 vs 0.739), and 0.014 on SCAIlZlTriplets (0.438 vs 0.424). Given Stella’s substantially larger footprint (1.5B parameters; 1536lZld embeddings) relative to Nomic (475M; 768lZld), selecting Nomic reduces index storage and retrieval cost by roughly 2 folds while maintaining comparable accuracy on our inlZldomain benchmark (SCAIlZlTriplets). Accordingly, we adopted NomiclZlV2lZlMOE as the dense embedding model for subsequent experiments.

**Figure 3.**
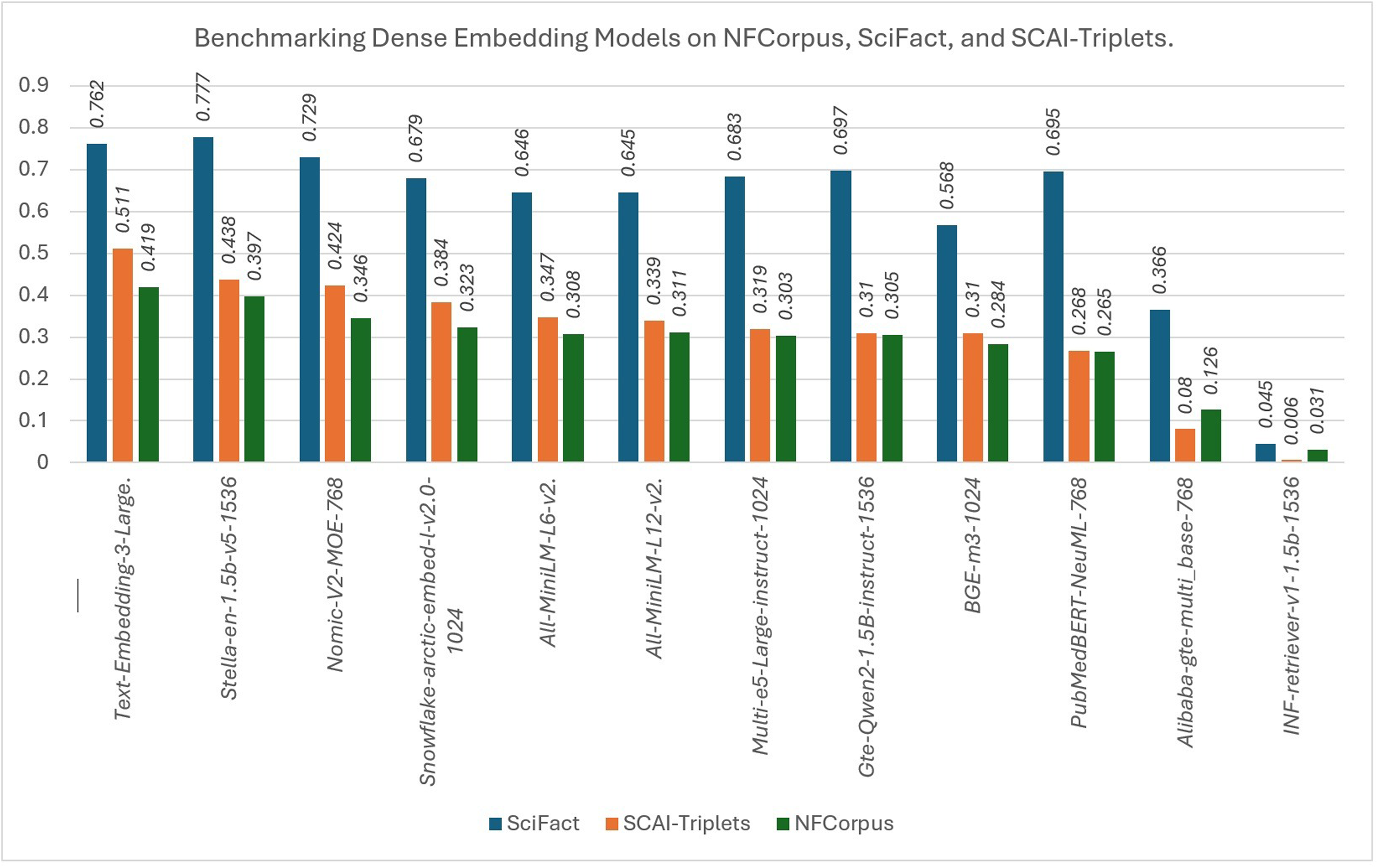
Benchmarking Dense Embedding Models on NFCorpus, SciFact, and SCAI-Triplets.

### 5.3 Hybrid Retrieval Enhances Open-Source Models Performance

After benchmarking the embedding models, we selected four top-performing open-source model families of varying parameter sizes —all-MiniLM variants (23M and 33M), Nomic-v2-MOE (475M), and Stella 1.5B (1.5B) — alongside with OpenAI’s text-embedding-3-large to further study the efficacy of hybrid versus dense retrieval. Averaged across NFCorpus, SciFact, and SCAIlZlTriplets, adding sparse embedding approach SPLADE to dense retrieval consistently improved or matched *nDCG*@10 for every model (Figure 4). Under Hybrid, OpenAI’s textlZlembeddinglZl3lZllarge achieved the highest mean *nDCG*@10 (0.570), while the strongest openlZlsource model (Stella 1.5BlZlv3lZl1536) reached 0.563, indicating near parity. NomiclZlv2lZlMOElZl768 attained 0.539, and MiniLM variants were ∼0.514. Absolute gains versus denselZlonly were largest for smaller models: Stella: +0.025 (0.538 → 0.563), Nomic: +0.039 (0.500 → 0.539), alllZlMiniLMlZlL6lZlv2: +0.082 (0.433 → 0.515), and alllZlMiniLMlZlL12lZlv2: +0.082 (0.432 → 0.514). These findings show that SPLADE benefits smaller/computelZlefficient models most, narrowing the quality gap between open-source and OpenAi’s model to ≤0.031–0.056 for midlZltier openlZlsource models and to ∼0.007 for the top openlZlsource model. When feasible, Hybrid retrieval is preferred, as it recovers ∼0.082 *nDCG*@10 for compact models and brings openlZlsource performance close to the proprietary baseline.

**Figure 4.**
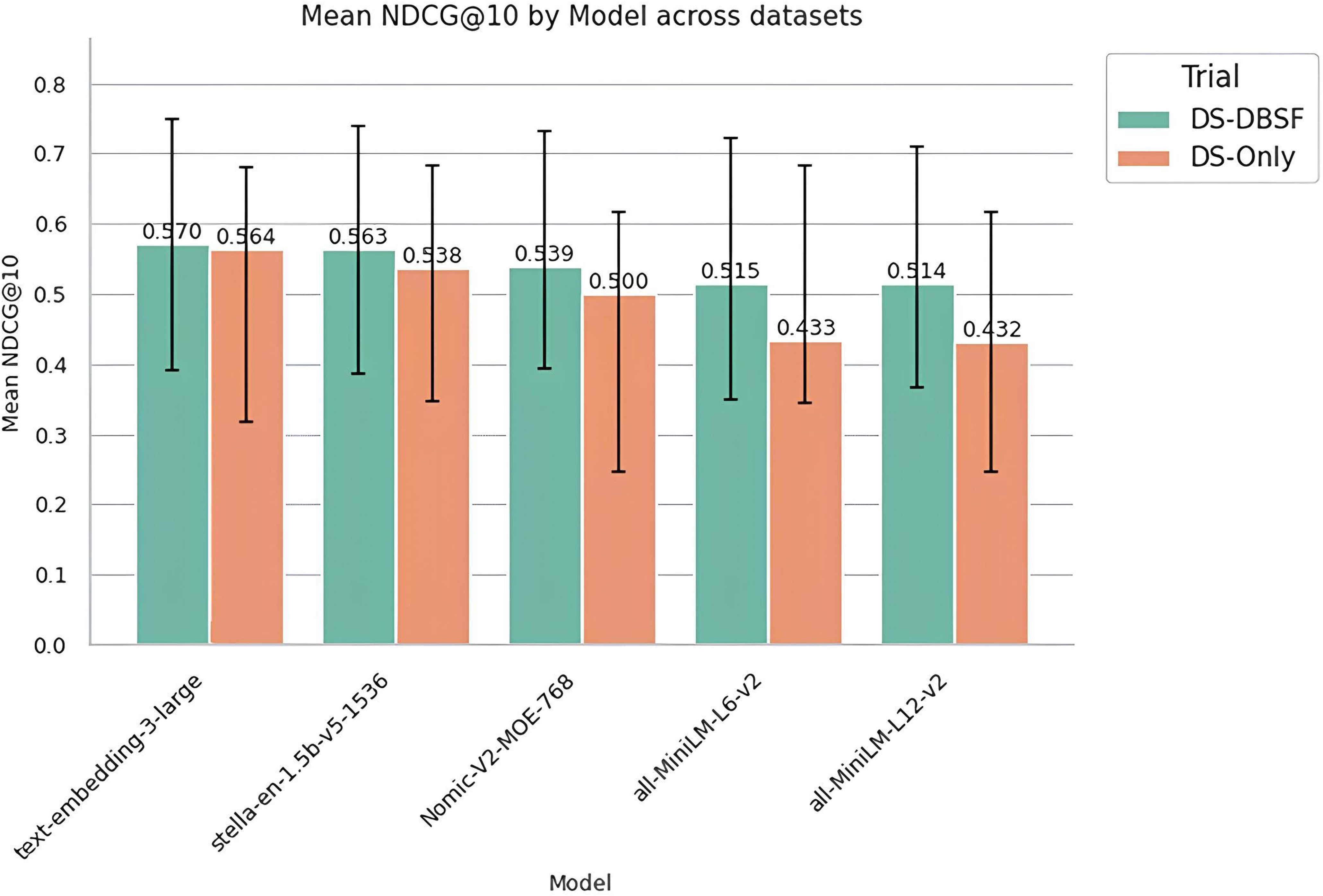
Comparison of Hybrid Retrieval using Dense and SPLADE Embeddings (DS-DBSF) versus DenselZlonly (DS-Only), averaged across nfcorpus, SciFact, and SCAIlZlTriplets.

### 4.4 Qwen 3 (32B) Demonstrates Superior Overall Quality

To identify the optimal backbone LLM for the agents, we benchmarked candidate models using NADKG-100, leveraging its compact size to minimize computational overhead during the initial selection phase. As shown in Tables 5 and 6, Qwen 3 (32B) outperformed the other models in both triplet-level and path-level assessments for NADKG-N100 enrichment, as evaluated using the OpenAI GPT-5-based evaluator. At the triplet level, it achieved biological validity 0.89, directionality validity 0.86, biomedical entity integrity 0.92, and triplet–evidence compatibility 0.75. Although Magistral (24B) generated the largest absolute number of triplets labeled valid, its per-criterion ratios were lower; only 0.56 of triplets were compatible with their evidence, and at the path level only 0.57 were causal/mechanistic, necessitating additional investigation and manual curation. DeepSeek-R1 (32B) showed solid biological validity (0.85) but lower triplet–evidence compatibility (0.68). Gemma 3 (27B) performed worst at the triplet level (compatibility 0.49; biological validity 0.66). At the path level, while Gemma 3 exceeded Qwen 3, DeepSeek-R1 and Magistral on most path-level metrics, model selection prioritized triplet-level quality—particularly biological validity and evidence because path coherence is not meaningful when constituent triplets lack support. Accordingly, Qwen 3 (32B) was selected as the backbone LLM for the framework.

**Table 5.**
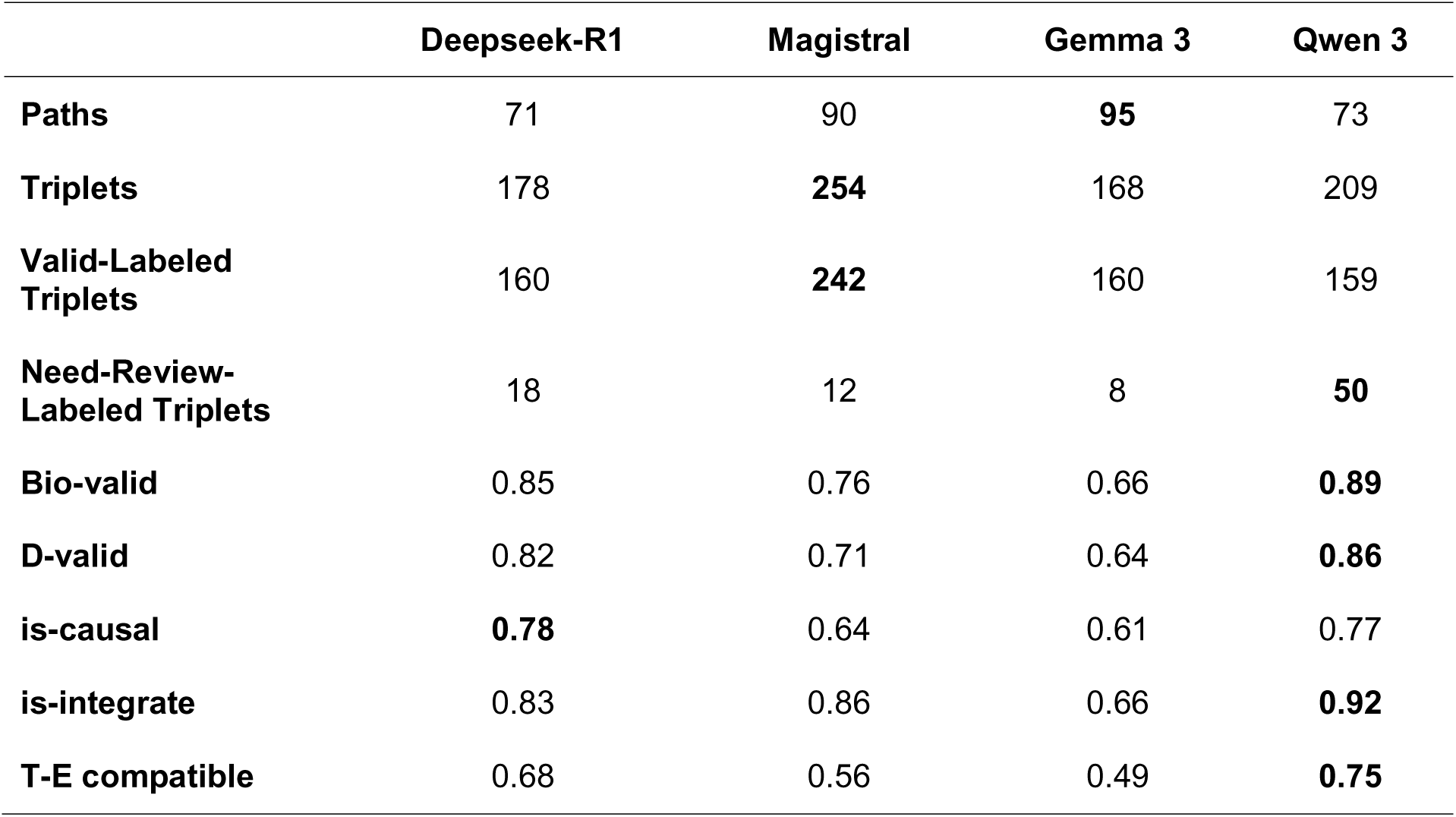
Triplet-level Evaluation of enrichment of NADKG-N100 for each LLM, evaluated by the OpenAI GPT-5–based evaluator (The highest value for each parameter is shown in bold).

**Table 6.**
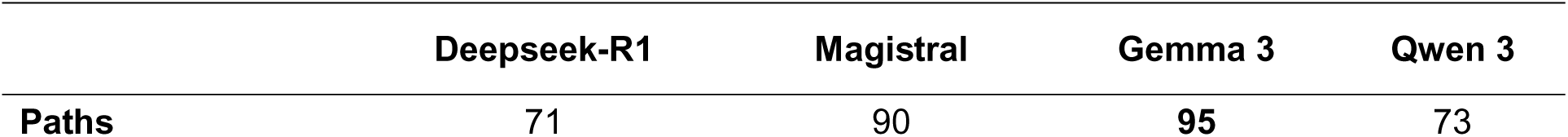
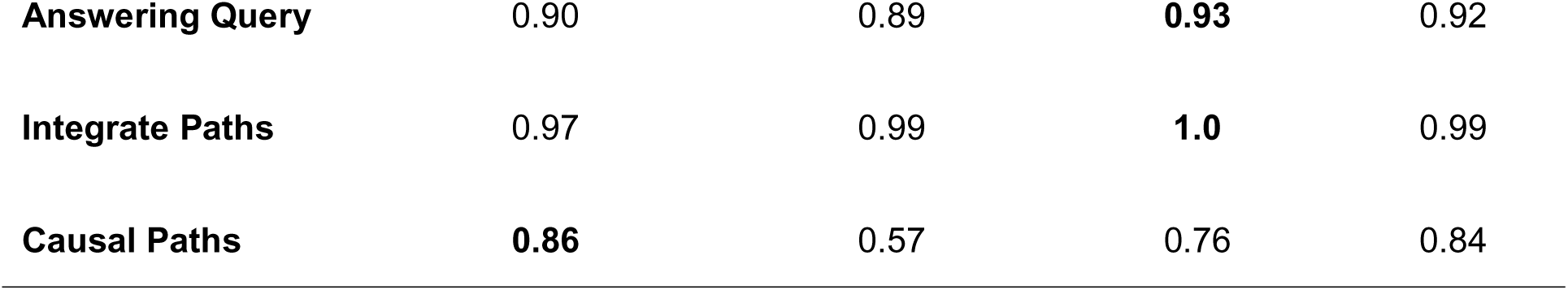
Path-level Evaluation of enrichment of NADKG-N100 for each LLM, evaluated by the OpenAI GPT-5–based evaluator (The highest value for each parameter is shown in bold).

### 5.5 Scaling Model Parameters Improves Reasoning and Coverage

After selecting Qwen 3 as the KGlZlOrchestra backbone, we compared the 14B, 32B, and 235B variants by enriching NADKG-N362 under identical prompting and evaluation settings, to examine scaling effects on enrichment quality and coverage. The 14B and 32B models were run on a single NVIDIA A100 (40 GB), and the 235B model on two H100 GPUs. As summarized in Tables 7 and 8, the 235B variant achieved the strongest tripletlZllevel performance (biological validity 0.97, directionality validity 0.96, causal/mechanistic coherence 0.83, entity integrity 0.95, and triplet–evidence compatibility 0.82) and produced 307 paths from 362 queries, confirming clear quality and coverage gains with increased capacity. Notably, the 14B model yielded slightly higher perlZltriplet quality than 32B but accepted fewer triplets and generated fewer paths (245/362). Despite the higher quality of generated paths compared to 32B and 235B, 14B showed limited reasoning abilities that hinder the understanding of more complicated paragraphs, which reduces recall and leaves more queries unanswered. Conversely, 32B extracted more valid hops and contributed more knowledge to the KG than 14B, at the cost of a small decline in perlZltriplet quality. On path-level performance, 14B achieved the highest pathways quality but extracted the least count of paths (245 paths) when compared to 32B (272 paths) and 235B (307 paths). Overall, KGlZlOrchestra remains robust under constrained computational power: 14B favors precision, 32B offers a strong coverage–quality tradelZloff, and 235B provides the best overall quality and path yield when resources permit.

**Table 7.**
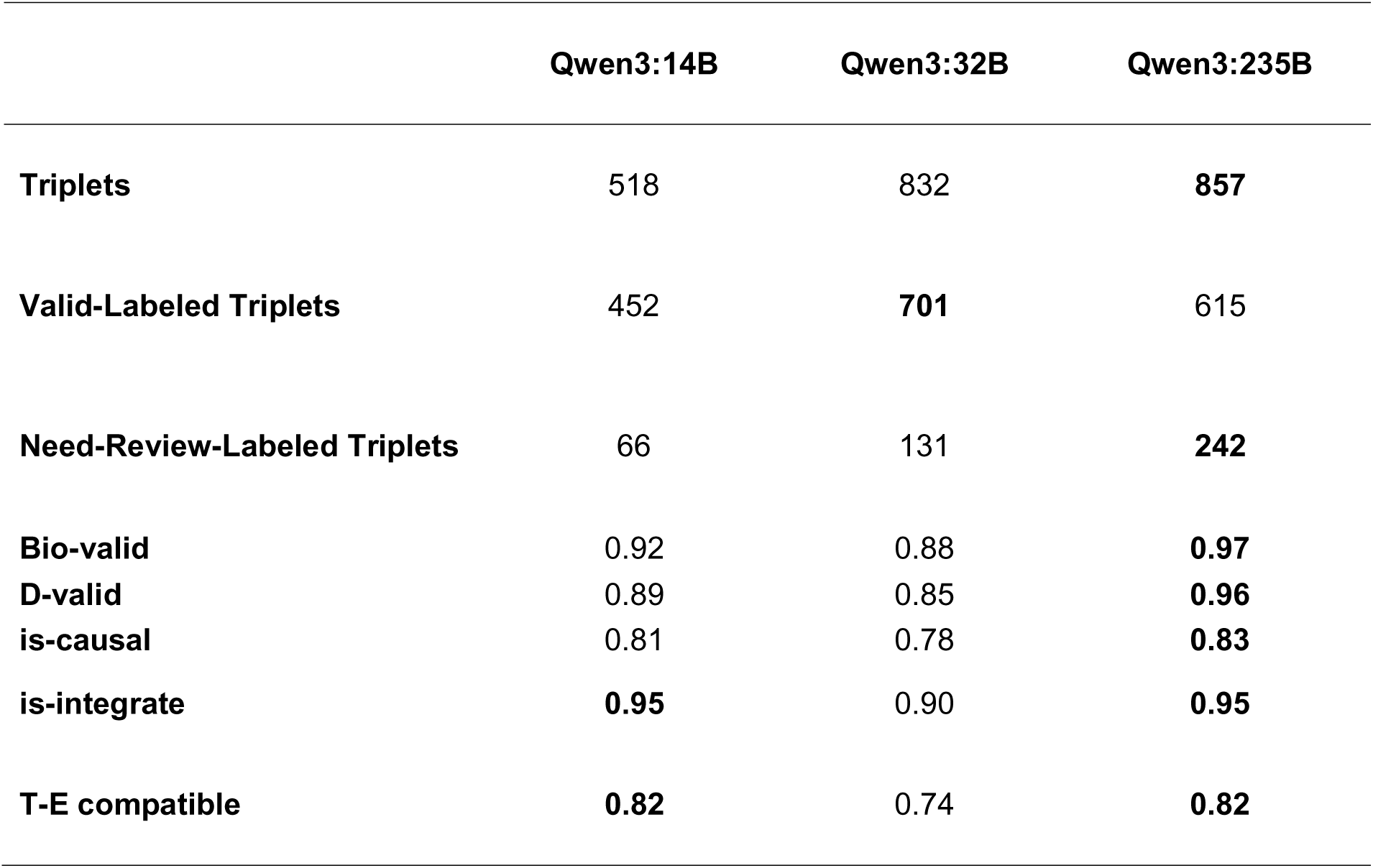
Triplets Evaluation Report generated from NADKG-N362 enrichment using different Qwen 3 versions (The highest value for each parameter is shown in bold).

**Table 8.**
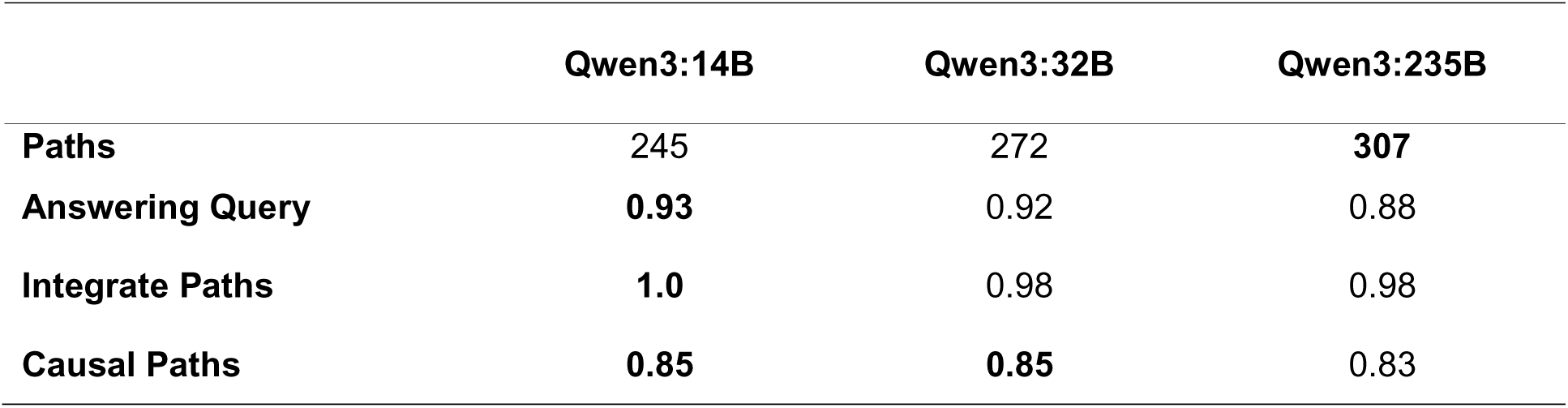
Paths Evaluation Report generated from NADKG-N362 enrichment using different Qwen 3 distilled versions (The highest value for each parameter is shown in bold).

### 5.6 Multi-Agent Validation Ensures High Triplet Quality

To quantify the marginal utility of each agent, given their added latency and compute cost, we ablated the query workflow and evaluated triplets produced at each stage of enriching NADKG-N362 using Qwen 3 (235B). As shown in Figure 5, Original triplets’ quality typically declines after aligning entity and relation types to the seed KG schema, whereas the Entity Matcher has negligible impact. Although running enrichment process without including the Schema Aligner in the workflow showed marginal triplet-level quality improvement (Figure 6), the final numbers of distinct entities and relation types were much higher in its absence (Table 9), underlining the essential role of the Schema Aligner in controlling ontology breadth and preventing type explosion, thereby preserving the graph’s analytic utility. Correspondingly, the Schema Aligner reduced the number of distinct head types from 78 to 49, tail types from 80 to 46, and relation types from 197 to 67 (Table 9). On the other hand, the Triplet Validation Team substantially improves triplet quality before ingestion, underscoring its critical role (Figure 5). It is important to notice that subsequent processing by the Triplet Validation Team increased distinct relation types to 76, reflecting selective introduction of semantically coherent relations. Overall, the ablation indicates minimal penalty from entity matching and significant quality gains from validation after the necessary schema alignment stage.

**Figure 5.**
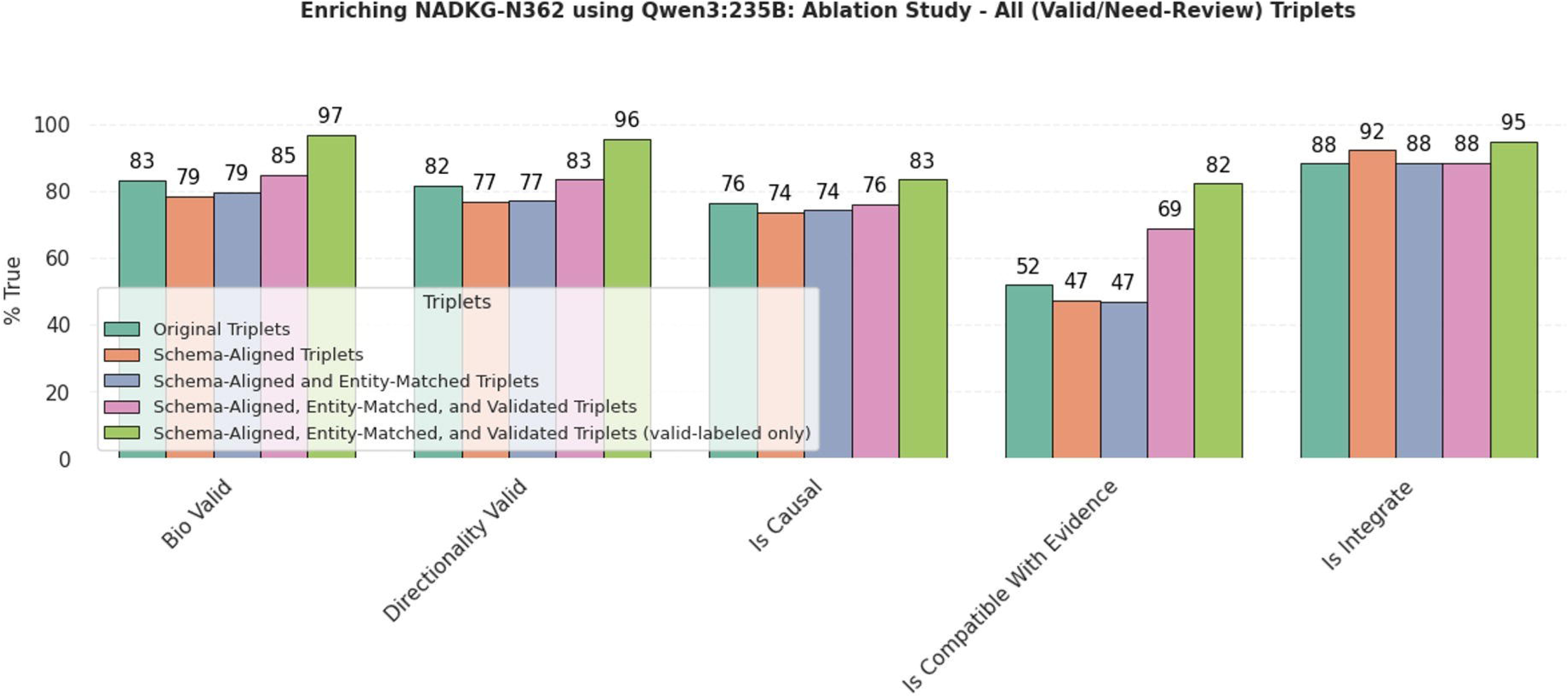
Ablation study of NADKG-N362 enrichment. Triplet quality was evaluated across four processing stages: (i) Original triplets, generated by the Path Builder; (ii) Schema-aligned triplets, produced by the Schema Aligner to conform to the Seed KG schema; (iii) Schema-aligned and entity-matched triplets, output of the Entity Matcher applied after schema alignment; and (iv) Schema-aligned, entity-matched, and validated triplets, produced by the Triplet Validation Team and labeled as “valid” or “need-review.”

**Figure 6.**
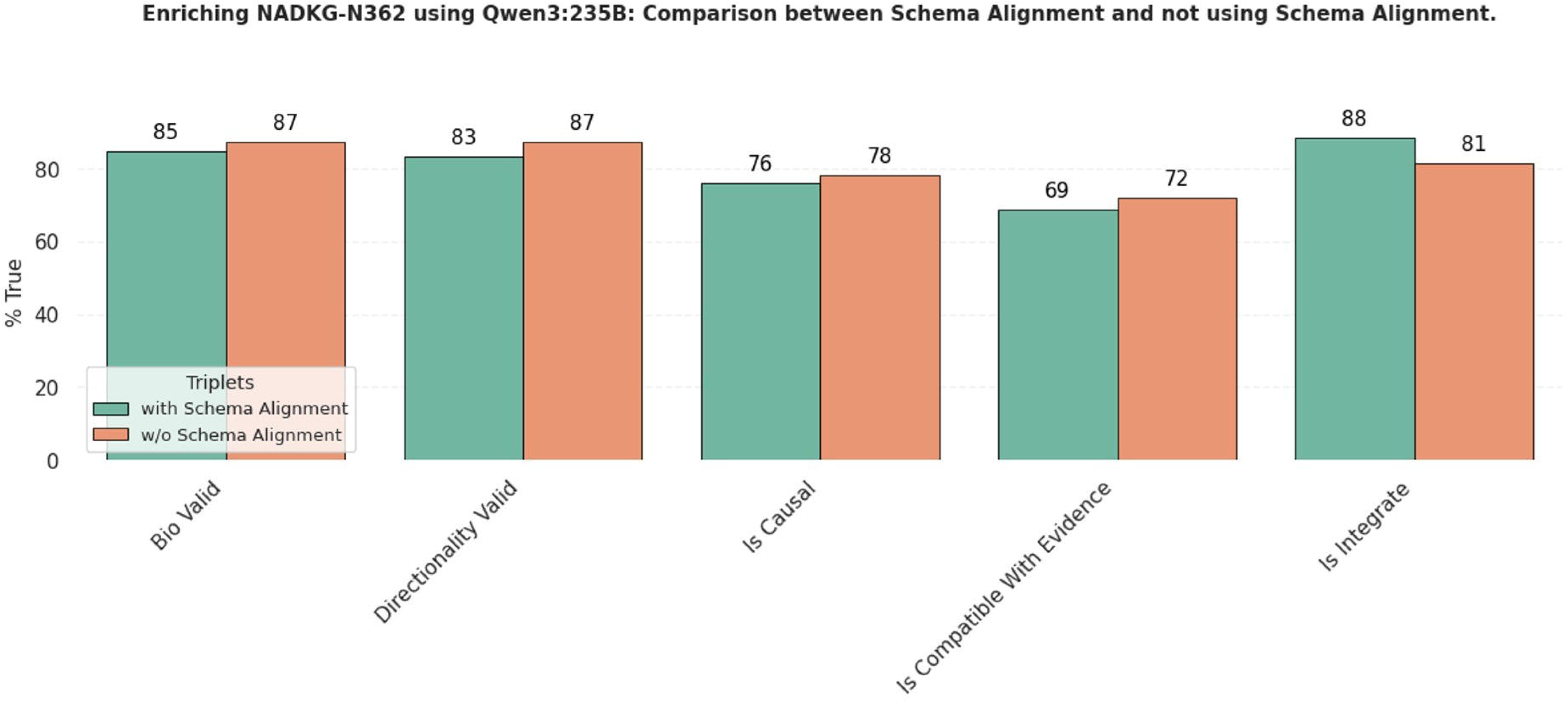
Ablation study of NADKG-N362 enrichment. Comparing triplet-level quality between using Schema Alignment versus not using Schema Alignment in KG-Orchestra’s workflow.

**Table 9.**
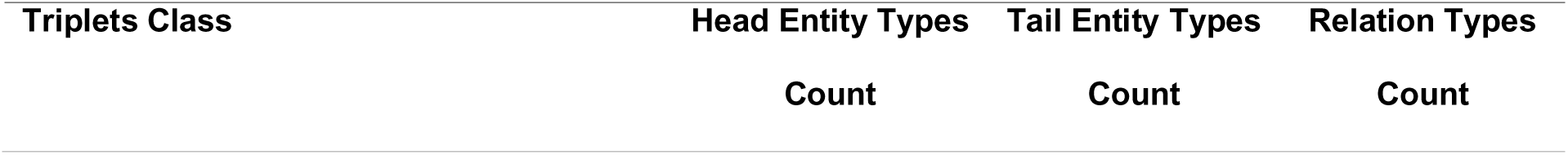
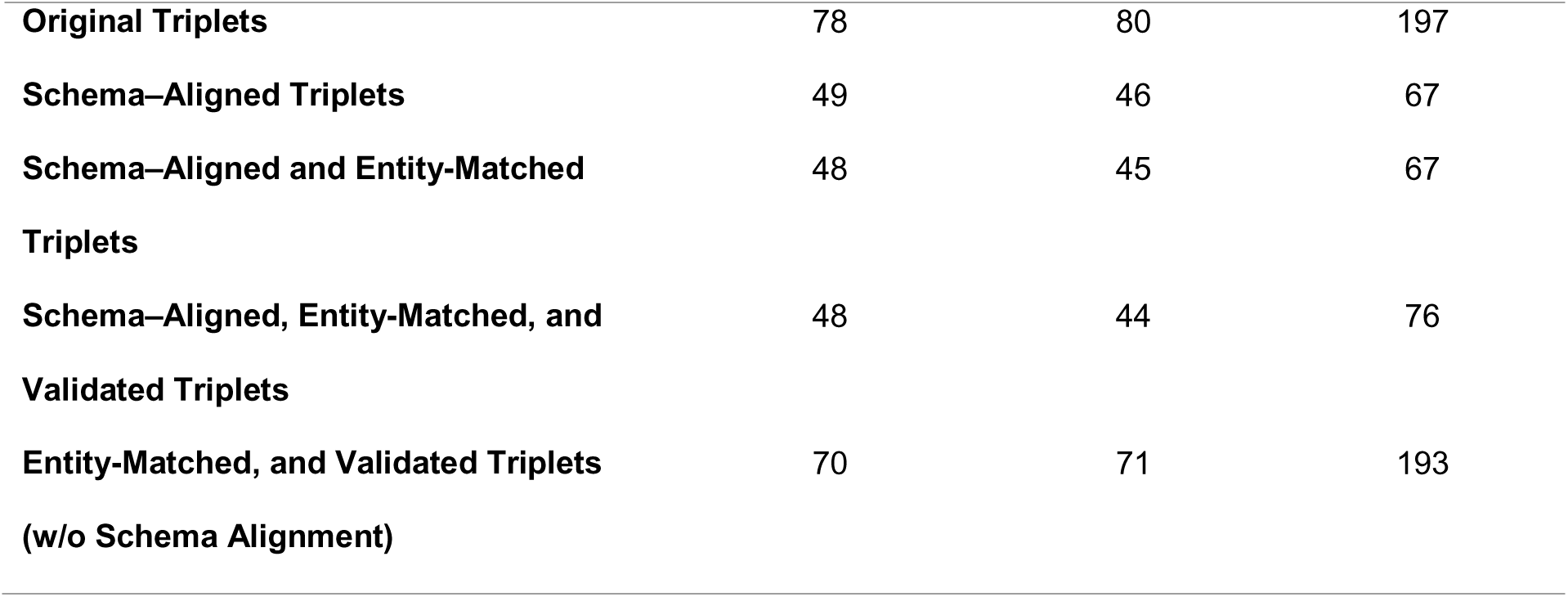
Distinct counts of head and tail entity types and relation types across successive stages of NADKG-N362 enrichment: (i) Original triplets, generated by the Path Builder; (ii) Schema-aligned triplets, produced by the Schema Aligner to conform to the seed KG schema; (iii) Schema-aligned and entity-matched triplets, output of the Entity Matcher following schema alignment; and (iv) Schema-aligned, entity-matched, and validated triplets, produced by the Triplet Validation Team and labeled as “valid” or “need-review.”

### 5.7 KG-Orchestra Expands Graph Coverage with High Biological Fidelity

We evaluated KG-Orchestra in an end-to-end setting by enriching the full ProPreSyn-GBA KG and NADKG and assessing a 20% random sample of the valid-labeled triples added to each enriched BKG using GPT-5, as automated evaluation, and manual evaluation by experts. On ProPreSyn-GBA (Figure 7; Table 10), KG-Orchestra added 1,037 nodes (approximately +141% relative to the seed) and 2,473 relations (approximately +182%), substantially expanding graph coverage. GPT-5-based automated evaluation of the triples sample indicated high quality: 93.1% biological validity, 77.2% relation causality, and 83.4% biomedical entity validity. Manual evaluation of the same triples sample yielded comparable estimates (Figure 7), corroborating the reliability of KG-Orchestra for biomedical KG enrichment. Comparable enrichment magnitudes and quality metrics were observed for NADKG (Figure 8; Table 10).

**Figure 7.**
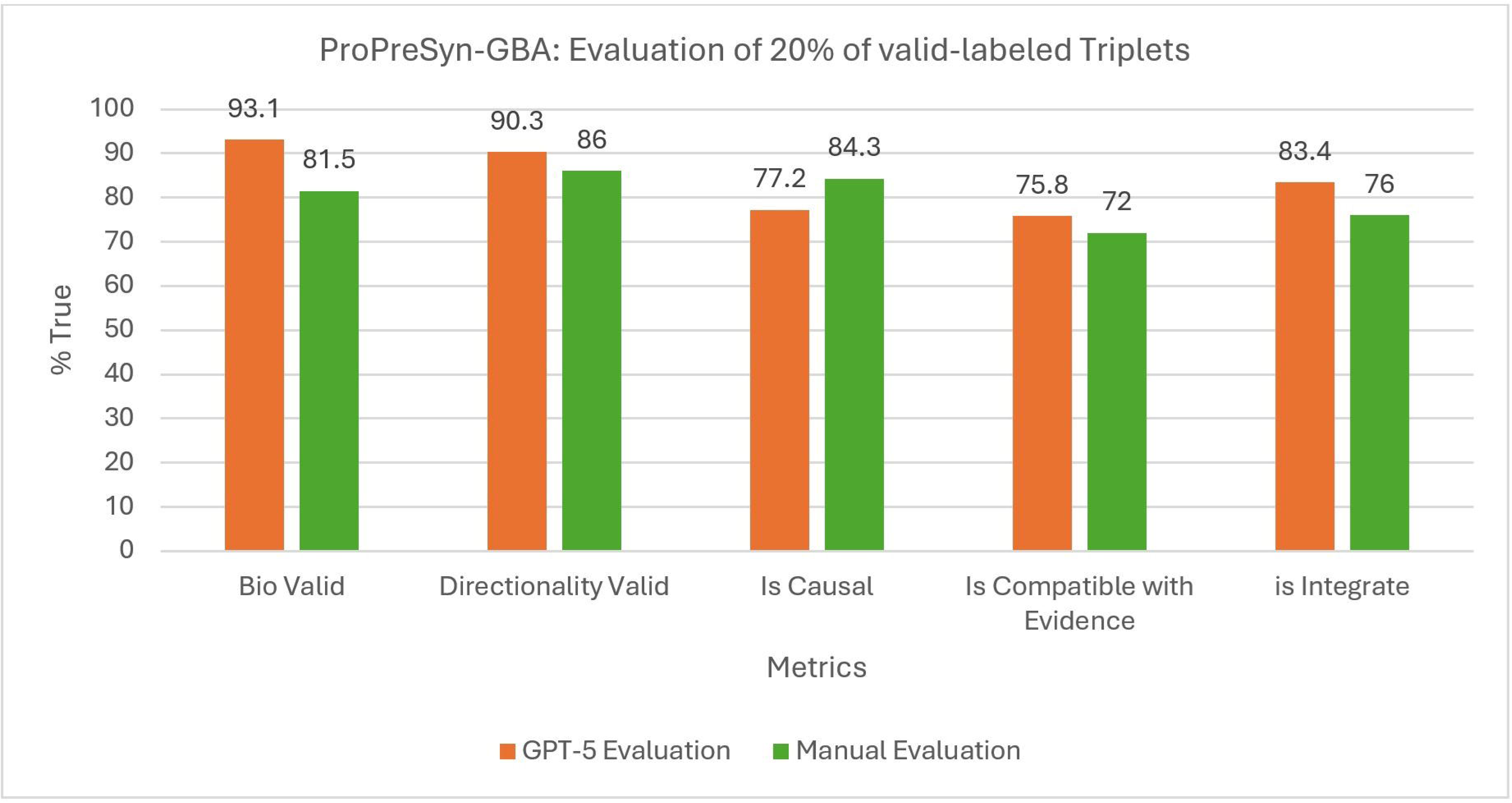
Evaluation of 20% of valid-labeled triplets generated during end-to-end enrichment of ProPreSyn-GBA seed knowledge graph.

**Figure 8.**
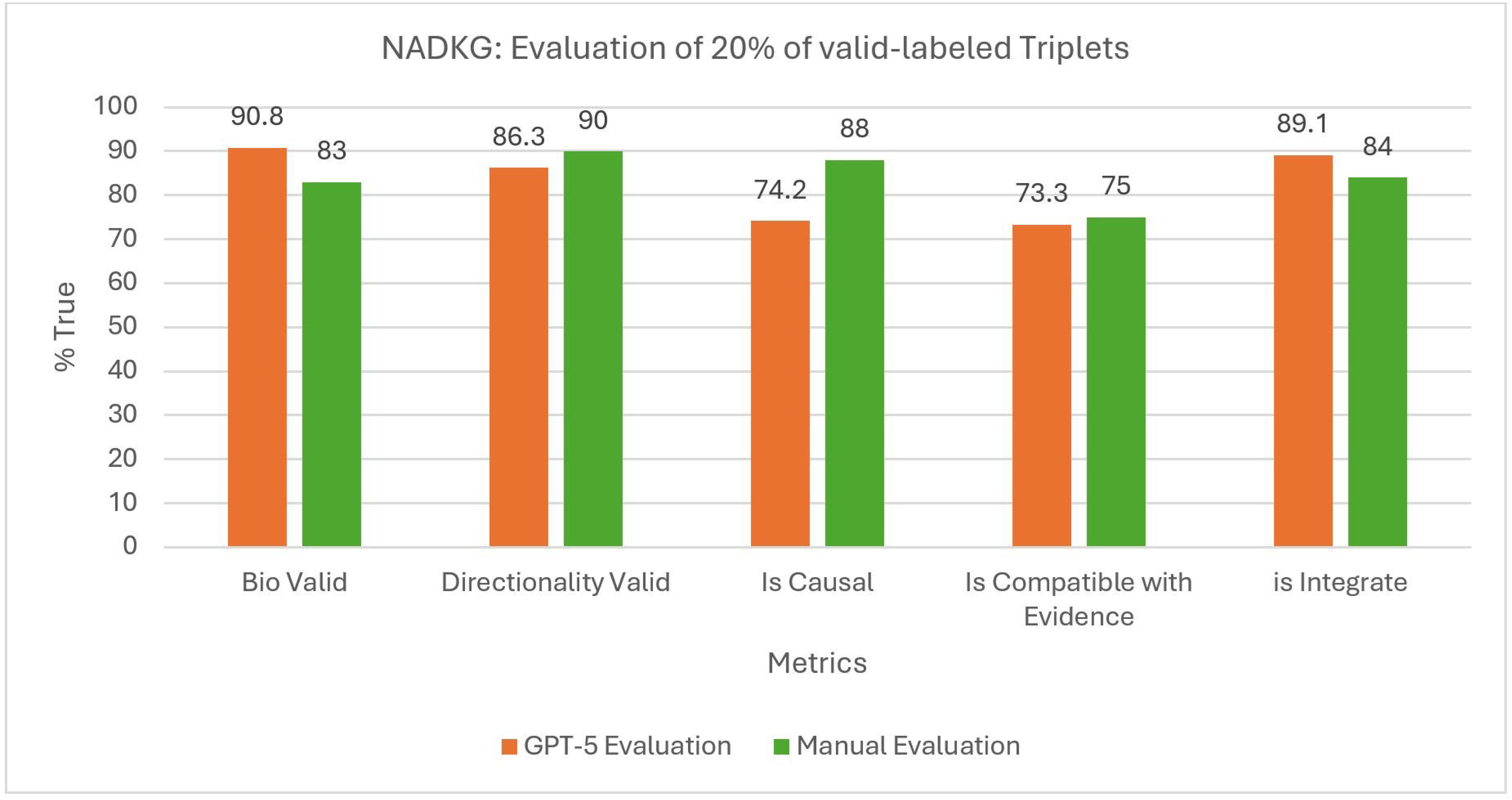
Evaluation of 20% of valid-labeled triplets generated during end-to-end enrichment of NADKG seed knowledge graph.

**Table 10.**
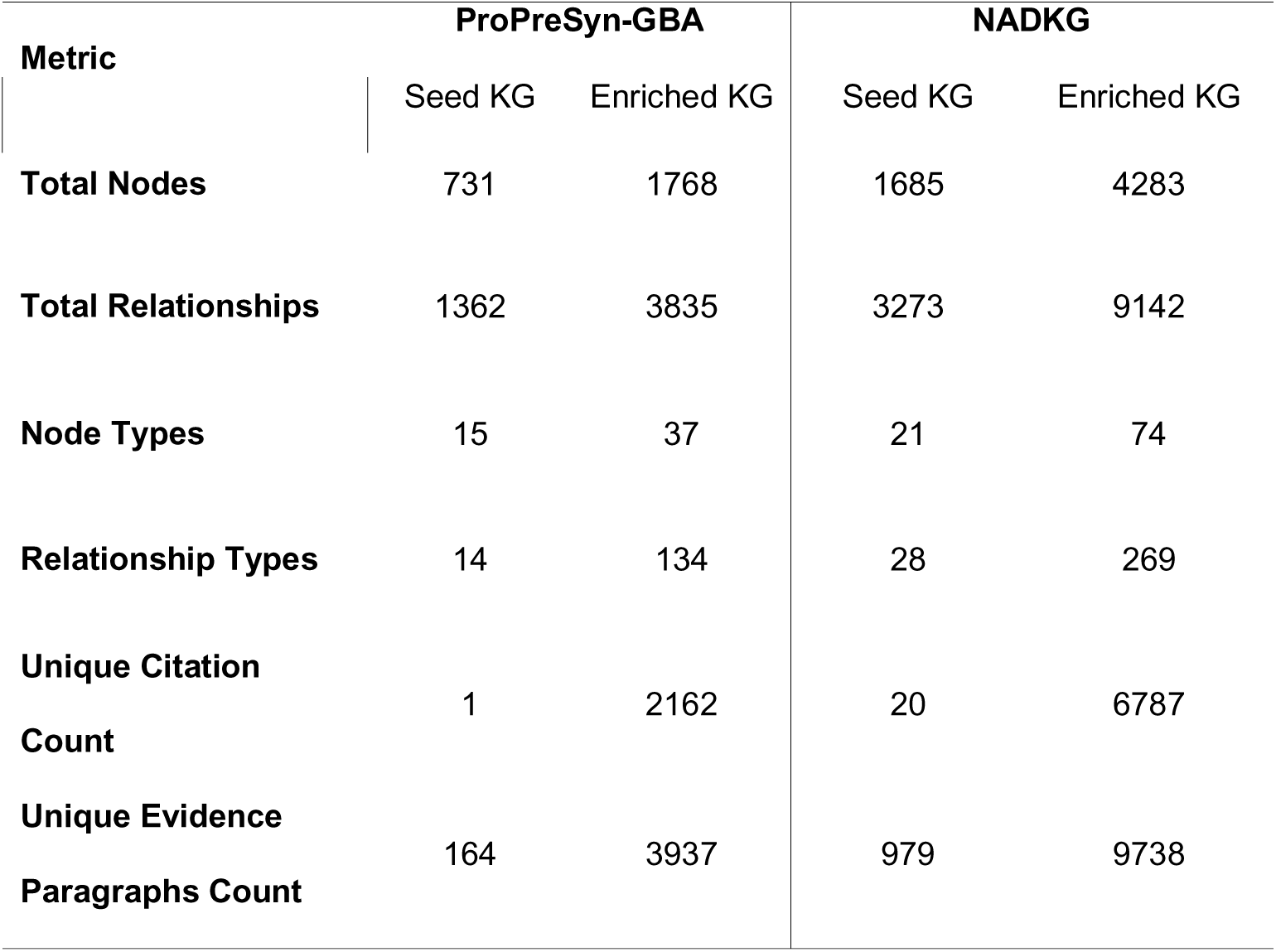
End-to-end enrichment of seed biomedical knowledge graphs by KGlZlOrchestra. For ProPreSynlZlGBA and NADKG, the table reports counts before (Seed KG) and after enrichment (Enriched KG) for: total nodes and relationships; number of node types and relationship types; unique citation count (distinct source articles); and unique evidence paragraph count (deduplicated supporting paragraphs). Enriched graphs include valid and need-review labeled triples.

As an illustration of the impact of enrichment, applying KG-Orchestra to NADKG recovered literature-supported, multi-hop paths linking Nelivaptan to Alzheimer’s disease. Figure 9 shows a subgraph capturing the hypothalamic–pituitary–adrenal (HPA) axis: Nelivaptan antagonizes the vasopressin V1B receptor (AVPR1B) [58], a stress-response node that increases corticotropin-releasing hormone (CRH) secretion [59,60], thereby promoting adrenocorticotropic hormone (ACTH) release [61] and consequent cortisol elevation [62]. Cortisol is positively associated with Alzheimer’s disease and neurofibrillary pathology, which in turn connects to Alzheimer’s disease [63,64]. Together, these relations form the chain: Nelivaptan —| AVPR1B → response to stress → CRH → ACTH → cortisol → neurofibrillary pathology → Alzheimer’s disease, demonstrating that enrichment via KG-Orchestra can recover valid, interpretable multi-hop paths integrating pharmacology, endocrine regulation, and neuropathology.

**Figure 9.**
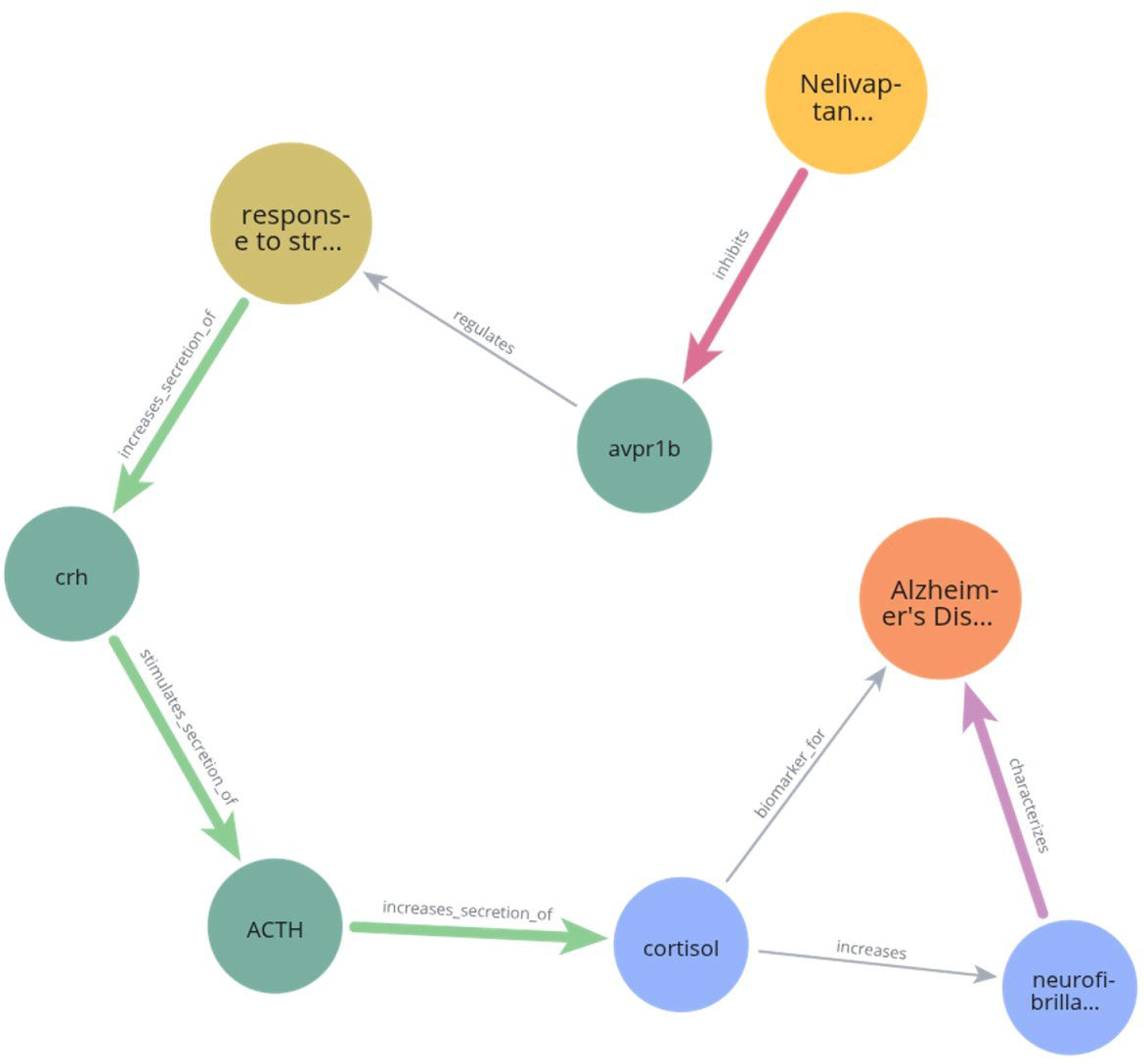
A path recovered by KG-Orchestra connecting Nelivaptan to Alzheimer’s disease within the enriched NADKG. Edges shown in grey represent relations present in the Seed NADKG prior to enrichment, whereas edges shown in red, green, and light purple correspond to KG-Orchestra–generated relations, each supported by evidence from the biomedical literature. “ACTH” node was introduced by KG-Orchestra during the enrichment process, while the other nodes were introduced by manual curators during the construction of seed NADKG.

### 5.8 The Framework Exhibits Robust Semantic Consistency

Across three independent executions of the KG-Orchestra pipeline on NADKG-N362, the framework produced highly consistent knowledge graph structures with only minor variability in graph composition. The number of extracted triplets remained stable across runs (820, 820, and 828 triplets, respectively), with comparable numbers of nodes (535–531–527) and relation types (64–60–66) (Table 11), indicating that the system reliably captures a similar semantic space even under the stochastic generation nature of LLMs. To quantify cross-run agreement at the level of extracted relations, we computed the symmetric Best-Match-average similarity between all pairs of triplet sets (see Methods). As shown in Figure 10, the resulting similarity matrix demonstrates strikingly high consistency, with pairwise scores ranging from 0.97 to 0.98 across all run comparisons. These values indicate that, for nearly every triplet in one run, a highly similar counterpart exists in the other runs, and vice versa. Collectively, these findings show that KG-Orchestra exhibits reproducible behavior, consistently converging on a stable set of semantic relations despite inherent randomness in LLM-driven extraction.

**Figure 10.**
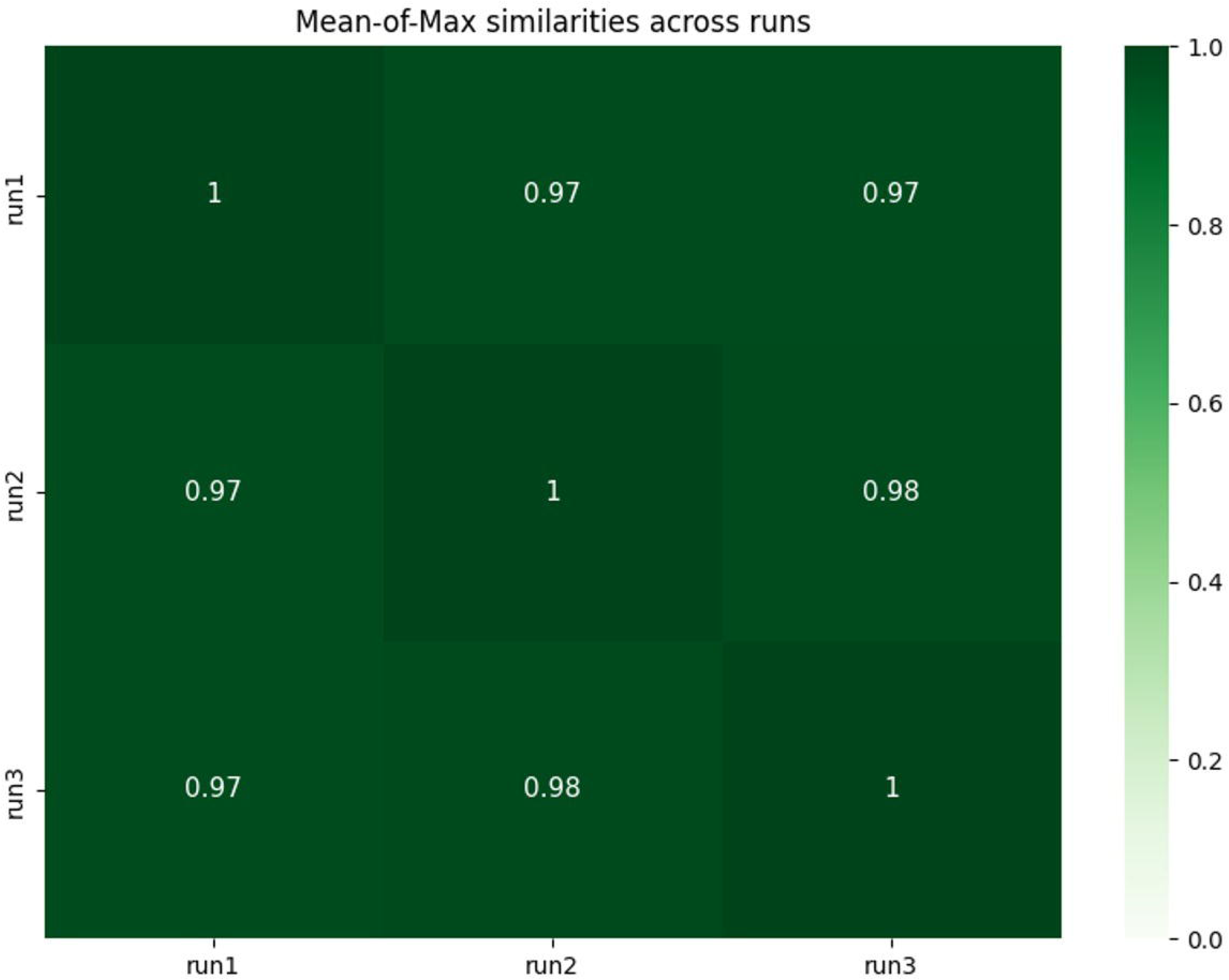
Similarity Matrix of Extracted Triplets Across Three Independent Runs Of KG-Orchestra on NADKG-N362.

**Table 11.**
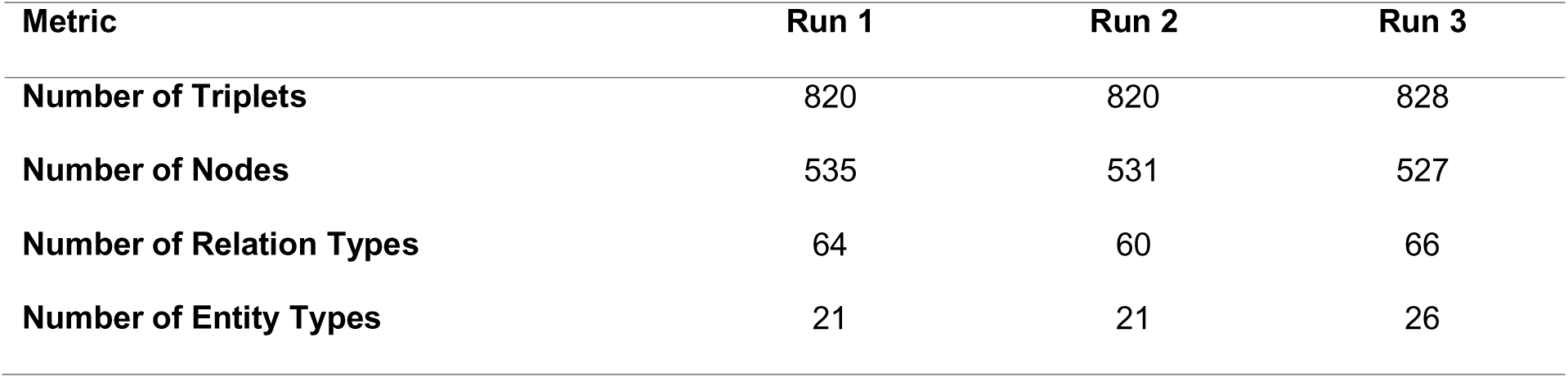
Extracted Triplets’ Characteristics of 3 independent runs of KG-Orchestra on NADKG-N362.

### 5.9 Seed KG Size Governs the Breadth of Retrieved Knowledge

We compared the extracted triplets after enriching NADKG-N100, NADKG-N362, and the full NADKG to assess how the size of the Seed KG influences downstream knowledge coverage, structural expansion, and triplet-level quality after enrichment. As expected, larger initial graphs produced substantially richer enriched KGs. The full NADKG contained 9142 triplets, 4283 nodes, 269 relation types, and 74 entity types, whereas NADKG-N362 expanded to 832 triplets, 549 nodes, 61 relation types, and 22 entity types. The smallest graph, NADKG-N100, yielded 209 triplets, 195 nodes, 35 relation types, and 15 entity types (Table 12). This monotonic increase across all structural metrics indicates that Seed KG size strongly governs the breadth of retrieved knowledge and the diversity of discovered semantic relations.

**Table 12.**
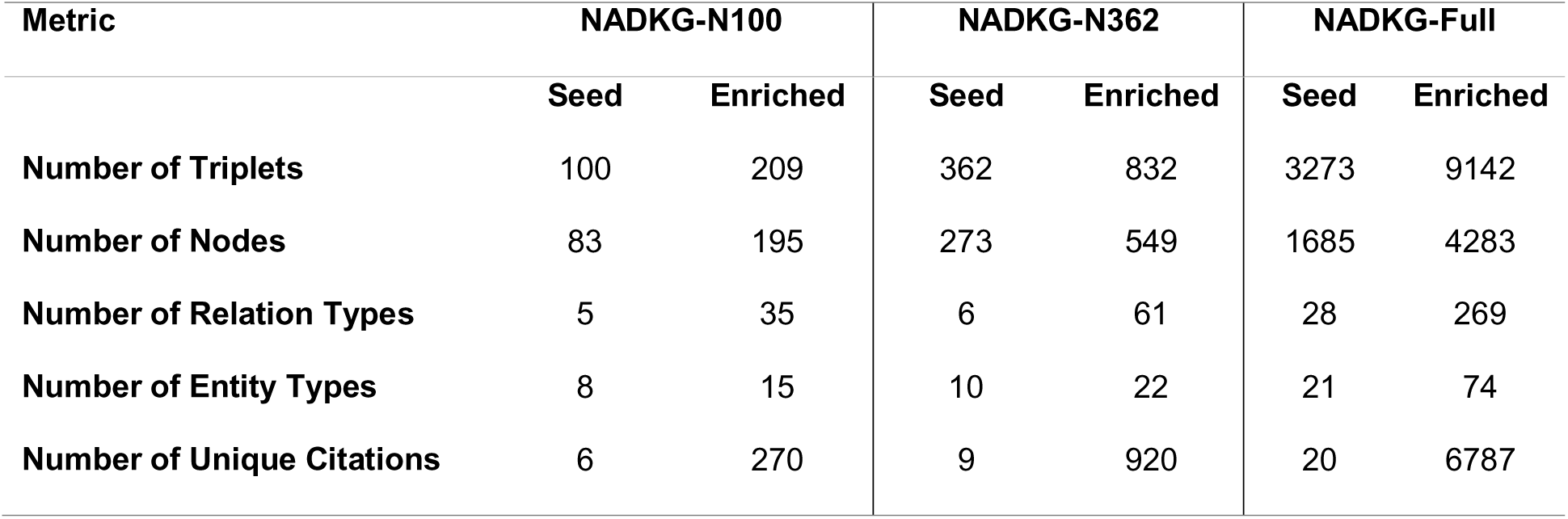
Comparison between Seed and Enriched NADKG-N100, NADKG-N362, and NADKG-Full Characteristics after enrichment.

Despite large differences in coverage, the triplet-level quality remained consistently high across all graph sizes (Figure 11). Biological validity ranged from 88–90.8%, directionality validity from 85–86.3%, and relation causality from 74.2–78%. Similarly, compatibility with evidence remained stable (73.3–75%), and entities’ integrity was high for all graphs (89.1–92%). These results demonstrate that while larger Seed KGs substantially increase the amount and diversity of recovered knowledge, they do not compromise triplet accuracy. Conversely, smaller KGs retain comparable semantic fidelity despite reduced coverage.

**Figure 11.**
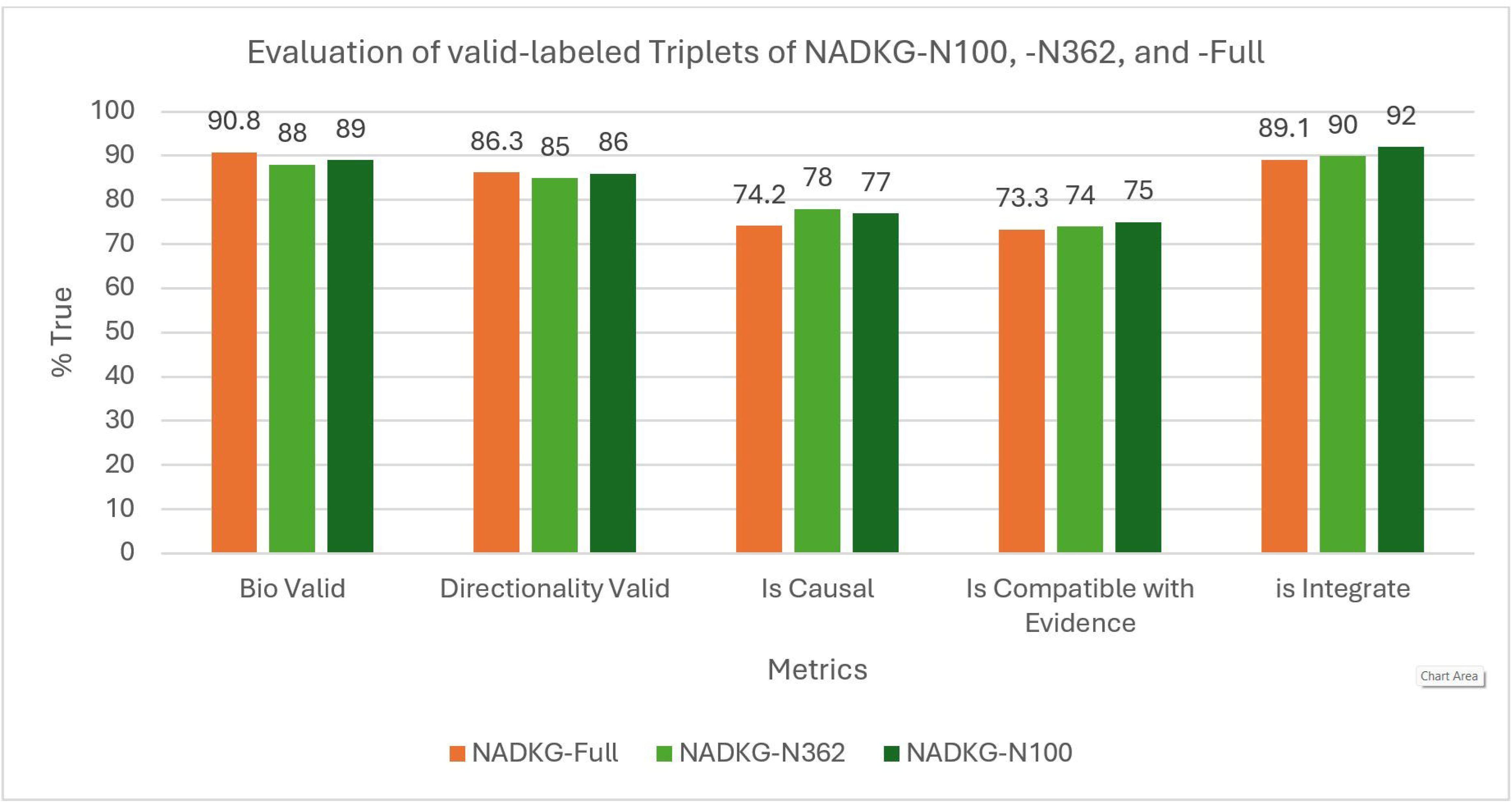
Comparison between NADKG-N100, NADKG-N362, and NADKG-Full Extracted Triplets’ Triplet-Level Quality after enrichment.

## 6. Discussion

The construction and enrichment of high-quality Biomedical Knowledge Graphs (BKGs) currently faces a fundamental paradox: while computational advances offer promising avenues for alleviating the burden on experts, the field still relies heavily on manual curation to ensure semantic precision. This persistent reliance stems from the fact that many automatically generated BKGs lack the quality and granular accuracy required for sensitive biomedical applications. However, the labor-intensive nature of manual curation creates a critical scalability issue, making it impossible to keep pace with the volume of new discoveries. This bottleneck prevents the timely enrichment of knowledge graphs with the most current and comprehensive information. To bridge this gap, there is a need for frameworks that prioritize high-fidelity extraction through a human-in-the-loop approach — leveraging automation to achieve the granularity and completeness required for specialized topics while maintaining the quality standards of expert-led verification. Furthermore, constructing and enriching a BKG from a limited number of biomedical articles inherently captures only a partial and potentially biased snapshot of the underlying scientific landscape. Biomedical knowledge is highly volatile – new discoveries, retractions, and revisions constantly reshape existing relationships among genes, diseases, drugs, and pathways. As a result, a BKG built from a static or restricted corpus risks omitting emerging concepts, misrepresenting causal links, and reinforcing publication or domain biases present in the source data. To mitigate these issues, an additional stage of graph validation and enrichment, without relying on a static or restricted corpus, is essential. The challenge extends beyond simply mitigating LLM hallucinations or verifying accuracy and biological coverage [65] to truly address the granularity of the captured information. A highly granular, context-specific biomedical knowledge graph enriched with causal relationships allows researchers to move beyond surface-level associations and instead interrogate the mechanistic fabric of biology. By capturing fine-grained entities — such as cell-type-specific gene functions, condition-dependent molecular interactions, or pathway dynamics under specific perturbations — and linking them through explicit causal edges, such a graph becomes a powerful engine for hypothesis generation. This level of detail supports truly discovery-driven science, allowing researchers to systematically uncover latent or previously unrecognized connections that are not obvious from isolated datasets or traditional literature mining. Ultimately, this enables the identification of non-canonical targets and empowers researchers to ask deeper, more precise questions about why biological phenomena occur, rather than just whether they correlate.

The overall findings of this study highlight how KG-Orchestra integrates retrieval, embedding selection, chunking strategies, and multi-agent reasoning into a coherent pipeline that performs reliably across a wide range of computational settings. Beginning with retrieval and chunking, KG-Orchestra enriched ProPreSyn-GBA and NADKG efficiently across diverse LLM configurations and model sizes, making the framework deployable under varied resource constraints. Within this context, 512-token-length-bounded hybrid chunking emerged as the highest-performing strategy, achieving the best NDCG@10 by preserving discourse-level semantics that sentence-only methods tend to lose. This advantage held across all benchmarking datasets, with the exception of the predominantly sentence-structured SCAI-Triplet corpus, where improvements were naturally marginal. These observations collectively anchor the importance of chunk granularity as the foundation upon which the remaining components of the pipeline build.

Building on the retrieval layer, model selection also shaped downstream performance. Nomic-v2-moe dense embedding model showed a good balance between size, computation cost, and retrieval quality. MixturelZloflZlExperts (MoE) models achieve superior performance in constrained computation environments by activating only a small subset of their total parameters for each input, effectively decoupling model capacity from inference cost [66]. This sparse activation allows MoE architectures to maintain the representational power of large models while significantly reducing memory usage and latency, enabling efficient deployment without sacrificing quality [67]. Additionally, the specialization of experts on different data subspaces enhances representational efficiency, enabling the model to allocate computational resources dynamically and achieve higher performance per unit of compute [68]. Within the broader KG-Orchestra workflow, this efficiency enables semantic retrieval to operate at scale and feed higher-quality representations into subsequent reasoning modules.

To further strengthen retrieval, combining Nomic-generated dense embeddings for semantic retrieval with SPLADE for lexical matching further increased *nDCG*@10 across all models. The implemented hybrid retrieval approach offers improved relevance because the two modalities complement each other’s strengths. Dense embeddings excel at capturing semantic similarity when queries and documents use different wording, while sparse lexical models excel at identifying exactlZlterm overlap and rare entity matches that dense models often miss [69]. By fusing both signals, the retrieval system attains higher overall effectiveness (e.g., better *nDCG*@10 scores) by balancing broad semantic recall with precise lexical precision, thereby improving the relevance of top-ranked evidence paragraphs.

Downstream of retrieval, Qwen 3 variants, as the backbone LLM for agents, consistently delivered superior performance, suggesting stronger context-sensitive reasoning. Increasing QwenlZl3 parameter counts yielded improved extraction from complex paragraphs and recovery of missing paths while maintaining tripletlZllevel quality. Qwen 3 models’ integration of a hybrid “thinking mode” and “nonlZlthinking mode” enables dynamic allocation of reasoning depth based on task complexity, thereby improving contextlZlsensitive reasoning without sacrificing latency [52].

In this multi-agent setting, dividing subtasks across specialized agents substantially improved reliability through cross-checking and self-correction, maintaining triplet quality throughout the workflow. Using multiple agents outperforms a singlelZlagent approach because each agent can specialize in a distinct subtask, allowing parallel exploration and verification of information. The ablation study shows that this division of labor enables crosslZlchecking and selflZlcorrection, reducing errors, and improving the consistency of final outputs. Consequently, multilZlagent systems achieve higher reliability and maintain quality across complex workflows compared with relying on a single agent’s sequential reasoning [70].

In line with these observations, our reproducibility analysis further confirms that KG-Orchestra produces highly consistent knowledge graphs across independent runs. Despite small fluctuations in the number of extracted triplets, nodes, and relation types, the semantic structure of the resulting graphs remains remarkably stable. The best-match-average similarity scores — ranging from 0.97 to 0.98 across all pairwise run comparisons — demonstrate that nearly every triplet generated in one run has a close semantic counterpart in the others. This high cross-run agreement indicates that the multi-agent pipeline not only improves intra-run quality through self-correction but also enhances inter-run reproducibility by converging on a robust and repeatable set of semantic relations. Together, these results show that KG-Orchestra’s agent-based architecture yields consistently reliable knowledge extraction even under stochastic generation conditions.

A related observation is that the size of the Seed KG influences the breadth of knowledge recovered during enrichment but has minimal impact on triplet-level accuracy, since the framework processes each triplet as an independent piece of information to be validated and enriched. Larger Seed KGs produced substantially richer graphs — containing more triplets, nodes, relation types, and entity types — indicating that initial graph scale primarily governs knowledge coverage. However, despite these large structural differences, triplet quality remained consistently high across NADKG-N100, NADKG-N362, and the full NADKG. Metrics such as biological validity, directionality accuracy, relations causality, compatibility with evidence, and entities integrity varied only minimally across graph sizes, demonstrating that KG-Orchestra maintains strong semantic fidelity even when operating with limited initial information. These findings highlight two key insights: the multi-agent architecture provides stable, accurate extraction regardless of Seed KG size, and larger graphs expand coverage without improving or degrading correctness. Collectively, this indicates that KG-Orchestra is both robust and flexible, capable of delivering reliable triplet-level reasoning in low-data settings while scaling effectively when richer prior knowledge is available.

Within this robust framework, KG-Orchestra supports four practical use cases that showcase its versatility. First, evidence-based biomedical knowledge graph completion: KG-Orchestra identifies and adds missing indirect paths between existing entities (nodes), linking each addition to citations from biomedical literature to improve graph coverage and support downstream applications that require comprehensive knowledge, such as novel drug target discovery. Second, introduction of new biomedical entities into seed graphs: the framework discovers and integrates paths connecting newly introduced entities to existing nodes, enabling rapid and scalable enrichment without extensive manual curation, with applicability across domains including linking new drugs to disease-specific graphs and informing drug repurposing. Third, construction of targeted biomedical subgraphs for specified entities: given a list of entities, KG-Orchestra automatically retrieves relevant evidence from large internal corpora and the web and assembles evidence-supported paths, reducing the need for manual article search and screening. Fourth, evidence enrichment and graph validation: for biomedical knowledge graphs with triplets lacking supporting citations, KG-Orchestra conducts targeted retrieval from curated corpora and trusted web sources, cross-validates each triplet against external evidence, and attaches detailed provenance (PubMed Central IDs/DOIs with linked supporting and contradicting excerpts), while flagging conflicts and uncertainty to enable transparent auditing and repeatable verification; this evidence-first process strengthens the evidential basis of graph assertions, enhances credibility and reproducibility, and improves reliability for downstream biomedical applications.

KG-Orchestra also presents several promising future applications, particularly in the context of automating ontology enrichment and construction. Its ability to identify literature-supported causal—direct or indirect—paths between biomedical entities provides a mechanism for uncovering mechanistic links that are frequently absent from manually curated resources. This is especially relevant for ontologies such as the Gene Ontology (GO) and for causal modeling frameworks like GO-CAM (Gene Ontology Causal Activity Models) [71], which build structured causal models from GO-defined activities. Currently, GO-CAM models require extensive manual curation to encode causal, regulatory, and functional relationships between biological activities, a process that is both time-consuming and difficult to scale as biomedical literature continues to expand [72]. KG-Orchestra’s automated extraction and inference of causal relations could substantially accelerate this workflow by proposing literature-grounded connections that enrich GO or assist in assembling GO-CAM models, ultimately reducing reliance on manual curation. Prior efforts in automating ontology construction have demonstrated that structured information can be systematically extracted from text to populate controlled vocabularies and relational structures, reinforcing the feasibility of replacing parts of the manual process with algorithmic support [73–75]. By applying this principle specifically to causal relations, KG-Orchestra offers a path toward more scalable, dynamically updated causal models within ontologies. In doing so, it could strengthen biological interpretability, support more rapid integration of new findings, and enable the continuous evolution of ontology-based knowledge representations in step with ongoing scientific discovery.

Despite the overall efficiency and robustness of KG-Orchestra, several limitations were observed. First, the Schema Aligner Agent occasionally fails to map extracted relation types to the broader causal or mechanistic polarity-aware relations defined within the seed KG schema, resulting in conceptual redundancy across entities and relationship types after enrichment. Second, the framework’s Evidence-Retrieval Pipeline is restricted to publicly accessible literature sources, constraining the breadth of incorporated knowledge and leading to incomplete coverage of several biomedical subdomains. Third, approximately 10–15% of generated triples were flagged as “need-review,” underscoring that a non-trivial portion of model-produced outputs still requires manual expert verification. Finally, because the retrieval module relies exclusively on dense and sparse embedding similarity, the system occasionally fails to surface the most semantically relevant paragraphs needed to satisfy a query. In such cases, the retrieval component may retrieve passages that are topically similar yet semantically irrelevant, while overlooking paragraphs containing essential causal or mechanistic evidence. These failures sometimes cause the pipeline to prematurely terminate and skip queries, highlighting the inherent limitations of embedding-only retrieval in capturing the inferential and mechanistic structure characteristic of biomedical literature.

Future development of KG-Orchestra will focus on enhancing both semantic fidelity and computational efficiency through three key integrations. First, we aim to incorporate semantic-operator frameworks like LOTUS [76] to replace standard vector similarity with reasoning-aware predicates, ensuring that retrieved evidence explicitly supports causal assertions rather than merely sharing topical keywords. Second, to capture the full complexity of biological mechanisms, we plan to embed a Biological Expression Language (BEL) translator, enabling the standardization of extracted claims into high-resolution, computable causal statements that support downstream reasoning [77]. Finally, to address the scalability constraints of deploying large-scale agents, we will evaluate emerging 1-bit Large Language Model architectures such as BitNet [78], which promise to drastically reduce inference latency and energy consumption while maintaining competitive reasoning performance. Collectively, these efforts will improve the robustness of schema alignment, broaden evidence coverage, and increase the reliability of automatically generated triples, ultimately minimizing the need for manual review.

## 7. Conclusion

In conclusion, KG-Orchestra demonstrates that effective enrichment of biomedical knowledge graphs hinges on the integration of curated domain corpora with context-aware retrieval strategies. By utilizing the seed knowledge graph as a schema and domain prior, the framework ensures granular yet low-noise expansion, efficiently adding missing entities and relations while validating existing assertions against diverse, up-to-date sources. The decomposition of workflows across specialized agents proves critical, enhancing reasoning capabilities through iterative self-correction and cross-agent validation, while the capture of stepwise reasoning summaries ensures process transparency and expedites curator-driven debugging. As an open-source solution, KG-Orchestra remains computationally flexible and deployable across a wide range of hardware—from single-laptop GPUs to high-performance clusters—by selecting the appropriate Qwen 3 model size. However, given the rapid velocity of biomedical discovery, sustained performance will require semantically aware acquisition pipelines and periodic corpus refreshes, as static keyword-based scraping is insufficient to prevent coverage gaps in the long term.

## 8. Author contributions

Ahmed Hossameldin Mohamed and Karim S. Shalaby conceived and designed the study. Ahmed Hossameldin Mohamed developed the framework, interpreted the results, and wrote the manuscript. Heval Atas Güvenilir participated in writing and proofread the manuscript. Abish Kaladharan conducted expert evaluation of the framework outputs. Alpha Tom Kodamullil acquired the funding and reviewed the manuscript. All authors have read and approved the final manuscript.

## 11. Data Availability

The data related to this article are available as an open-source Python package at https://github.com/Fraunhofer-SCAI-Applied-Semantics/KG-Orchestra under the Apache 2.0 License.

## 13. Glossary

Ablation Study: A scientific procedure used to determine the contribution of individual components of an AI system by removing them one at a time and measuring the resulting impact on performance.
AI Agent: An autonomous computational entity driven by a Large Language Model that is designed to perform a specific, specialized task—such as text retrieval, schema alignment, or fact-checking—within a larger workflow.
Biomedical Knowledge Graph (BKG): A structured network representation of biological knowledge where nodes represent entities (e.g., genes, drugs, diseases) and edges represent the semantic or causal relationships between them.
Embedding (Dense vs. Sparse): A mathematical representation of text as a numerical vector. **Dense embeddings** capture semantic meaning and context, while **sparse embeddings** (like SPLADE) focus on specific keyword matching and lexical importance.
Evidence: Within this framework, a specific paragraph-level textual excerpt retrieved from scientific literature that provides direct support for, or contradiction of, a proposed knowledge assertion.
Hybrid Retrieval: A search methodology that combines both dense and sparse embeddings to improve the accuracy and relevance of retrieved documents by balancing semantic intent with keyword precision.
Knowledge Graph Enrichment: The process of expanding an existing knowledge graph by adding new entities, discovering novel relationships, or validating existing assertions using external data sources.
Large Language Model (LLM): A type of artificial intelligence trained on massive datasets of text to understand, generate, and reason with natural language (e.g., Qwen, GPT-4, DeepSeek).
Multi-Agent System (MAS): An architectural framework where multiple specialized AI agents collaborate to solve complex tasks by decomposing them into smaller, manageable sub-processes.
NDCG (Normalized Discounted Cumulative Gain): A standard metric used in information retrieval to measure the quality of ranking; it rewards systems for placing the most relevant results at the top of a list.
Ontology: A formal and standardized set of definitions, categories, and properties that dictate how entities and relationships should be structured within a specific domain.
Provenance: The documented history of a data point, including its original source (e.g., DOI or PubMed ID) and the computational steps taken to extract and validate it.
Retrieval-Augmented Generation (RAG): A technique that enhances LLM performance by retrieving relevant documents from an external corpus and providing them to the model as context before generating a response.
Schema Alignment: The process of mapping newly extracted entities and relationships to the predefined structure and vocabulary of an existing knowledge graph to ensure consistency.
Seed Knowledge Graph: An initial, often smaller, knowledge graph that serves as the foundation and structural guide for further enrichment and expansion.
Triplet: The fundamental unit of a knowledge graph, consisting of three parts: a Head (subject), a Relation (predicate), and a Tail (object), forming a statement such as (*Drug A* — *treats* — *Disease B*).

